# Polycomb Repressive Complex 2 shields naïve human pluripotent cells from trophectoderm differentiation

**DOI:** 10.1101/2021.08.21.457215

**Authors:** Banushree Kumar, Carmen Navarro, Nerges Winblad, John P Schell, Cheng Zhao, Fredrik Lanner, Simon J Elsässer

## Abstract

The first lineage choice made in human embryo development separates trophectoderm from the inner cell mass, which proceeds to form the pluripotent epiblast and primitive endoderm. Trophectoderm on the other hand gives rise to the placenta. Naïve pluripotent stem cells are derived from the pluripotent epiblast of the blastocyst and offer possibilities to explore how lineage integrity is maintained. Here, we discover that Polycomb repressive complex 2 (PRC2) restricts an intrinsic capacity of naïve pluripotent stem cells to give rise to trophectoderm. Through quantitative epigenome profiling, we find that broad histone H3 lysine 27 trimethylation (H3K27me3) hypermethylation is a common feature of naïve pluripotency across species. We define a previously unappreciated, naïve-specific set of bivalent promoters, featuring PRC2-mediated H3K27me3 concomitant with H3K4me3. Naïve bivalency maintains key trophectoderm transcription factors in a transcriptionally poised state that is resolved to an active state upon depletion of H3K27me3 via inhibition of the enzymatic subunits of PRC2, EZH1/2. Conversely, primed human embryonic stem cells cannot be driven towards trophectoderm development via PRC2 inhibition. While naïve and primed hESCs share the majority of bivalent promoters, PRC2 contributes to the repression of largely non-overlapping subsets of these promoters in each state, hence H3K27me3-mediated repression provides a highly adaptive mechanism to restrict lineage potential during early human development.

## INTRODUCTION

Human embryonic stem cells (hESC) play a pivotal role in our pursuit of understanding mammalian development. *In utero,* the developmental cascade is triggered with the specification of the inner cell mass (ICM) and the trophectoderm (TE), the outer envelope - trophoblast - of the pre-implantation blastocyst. Upon implantation, the trophoblast gives rise to placental tissues while the ICM progresses via the epiblast stage to form the fetus. hESCs derived from the ICM of the developing blastocysts ^1^ have the capacity to self-renew and differentiate into any of the three (ectoderm, mesoderm and endoderm) germ lineages, a feature defined as pluripotency. Intriguing data ^2–9^ also suggest that hESCs, in contrast to mouse embryonic stem cells (mESCs), can be induced to differentiate into extraembryonic lineages such as trophectoderm cells, a capacity normally restricted to totipotent stem cells. The mechanisms restricting epiblast cells from entering trophectoderm lineage in the developing blastocyst remain elusive.

To align the pluripotency continuum during human embryogenesis with *in vitro* cultured cells, initial parallels were drawn between hESCs and the relatively well-characterized mESC systems. Conventional hESCs (grown with serum and fibroblast growth factor 2) resembled the post-implantation epiblast and its features clustered closer to the murine epiblast stem cells (EpiSCs) ^10, 11^, emanating the name: primed hESCs ^12^. To stabilize hESCs in the pre-implantation-like state, referred to as ’naïve’ state throughout this study, numerous protocols have been described ^13–18^. Though these conditions recapitulated by large the gene expression, X chromosome reactivation and DNA methylation patterns observed in the pre-implantation ICM *in vivo*, the diversity in signaling pathways targeted to obtain naïvety necessitated more definitive markers to define naïve hESC. In an effort to address this, several genomic, transcriptomic and proteomic analyses have been conducted to further characterize the cellular states and understand the molecular mechanisms governing them ^19–25^.

Epigenetic signatures, specifically histone post translational modifications (hPTMs) have emerged as a criterion to evaluate cell state transitions. Analogous to mESCs, in hESCs the conversion from primed to naïve, is accompanied by changes in deposition of repressive marks H3K27me3 and H2AK119Ub by the polycomb repressive complexes (PRC2 and PRC1 respectively) ^24, 26, 27^. In addition, bivalency, presence of both the repressive mark H3K27me3 and active mark H3K4me3 on developmentally regulated genes, has also been reported to be reduced in mESC ^28–30^. However, not much effort has been directed towards exploring a possible role of these epigenetic signatures in preventing differentiation, owing partly to the fact that PRC2 is dispensable for maintaining pluripotency in hESCs ^31, 32^.

Using MINUTE-ChIP ^33^, a quantitative ChIP-seq method, we profile changes in specific histone modifications between naïve and primed hESCs. We observe an increased pervasive deposition of H3K27me3 and H2AUb throughout the genome of naïve hESCs. In addition, with quantitative profiles of H3K4me3 we discover specific trends at bivalent promoters in different pluripotent states. We confirm that the X-chromosome is decorated with H3K27me3 in the naïve state albeit with no significant effect on gene expression. However, comparing epigenetic and transcriptional changes following treatment with an EZH2 (catalytic subunit of PRC2) inhibitor resulted in identification of a new set of naïve specific PRC2 targets that overlap with trophectoderm and extraembryonic tissue markers. Interestingly, removal of H3K27me3 is also accompanied by rewiring of H3K4me3 levels. Taken together, we elucidate a novel role of PRC2 in maintaining naïve pluripotency in humans.

## RESULTS

### Quantitative maps reveal genome-wide H3K27me3 hypermethylation and specific gains and losses of H3K27me3 at bivalent promoters in naïve human pluripotent stem cells

To elucidate the function of promoter bivalency in naïve and primed human pluripotency, we designed a quantitative ChIP-seq experiment for which H9 female hESCs were maintained in naïve or primed culture conditions, and treated with and without EZH2i (EPZ-6438; ^34^) for 7 days (Fig. 1a). We profiled three histone modifications associated with bivalent genes, H3K4me3, H3K27me3 and H2Aub. Triplicate samples of each condition were combined in a MINUTE-ChIP experiment (Extended Data Fig. 1a) ^33^. Phenotypically, EZH2i-treated naïve and primed hESCs maintained their expected morphology (Extended Data Fig. 2a).

**Fig. 1.**
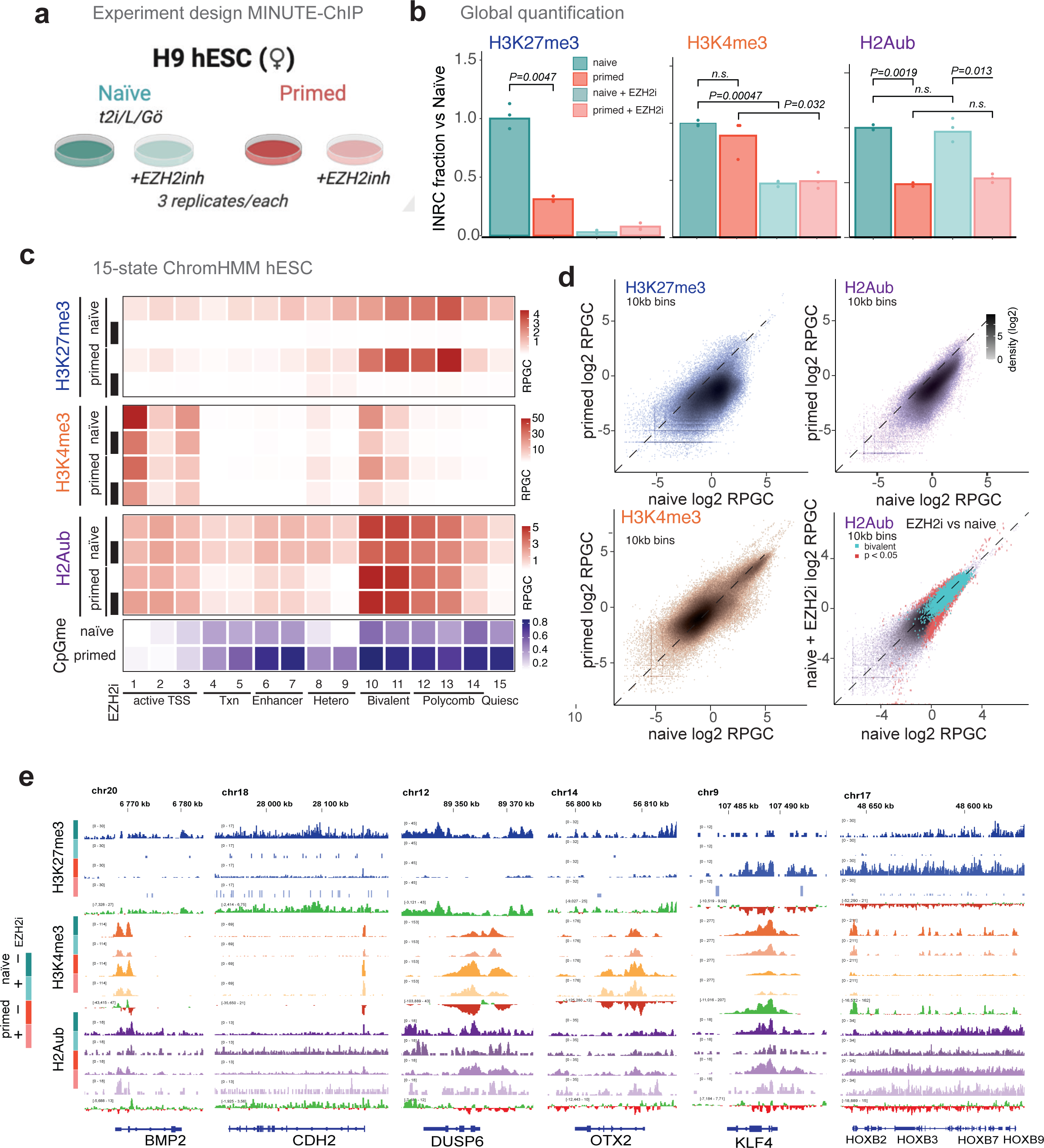
High levels of H3K27me3 and H2Aub broadly cover the naïve pluripotent genome. **a** Experimental design of multiplexed quantitative ChIP (MINUTE-ChIP) experiment comparing hESCs in naïve and primed culture conditions. In addition, both naïve and primed cultures were treated with EZH2 inhibitor EPZ-6438 (EZH2i) for 7 days. Three replicates of each condition were barcoded and combined into a single MINUTE-ChIP pool from which H3K27me3, H3K4me3 and H2AK119ub (H2Aub) ChIPs were performed. See Extended Data Fig. 1a for a scheme of the MINUTE-ChIP workflow. **b** Global genome-wide levels of H3K27me3, H3K4me3 and H2Aub as determined by input-normalized total read counts (INRC) in naïve or primed hESC, cultured with or without EZH2 inhibitor. P values of pairwise comparisons (Student’s t test) are given. Tracks for subsequent analysis are scaled according to the INRC values as reads per genome coverage (RPGC), with naïve serving as a reference scaled to 1x genome coverage (global average equals to 1 RPGC) **c** Histone H3K27me3, H3K4me3 and H2Aub levels by chromatin state. RPGC of combined replicates are shown. **d** Genome-wide analysis of H3K27me3, H3K4me3 and H2Aub levels by 10kb bins, comparing naïve and primed hESC. Log2-transformed RPGC of combined replicates are shown. Bottom right: analysis of genome-wide 10kb bins comparing H2Aub levels in naïve hESC, with or without EZH2i treatment. Differentially occupied regions are highlighted in red (DESeq2 p.adj < 0.05). Bins overlapping with annotated bivalent promoters ^62^ are marked in turquoise. **e** Genome browser examples of genomic regions with differential occupancy of H3K27me3, H3K4me3 and/or H2Aub. Shown is the combined signal of three replicates, gain/loss tracks comparing naïve and primed signal. Tracks from the same histone modification are shown on the same RPGC scale.

**Fig. 2.**
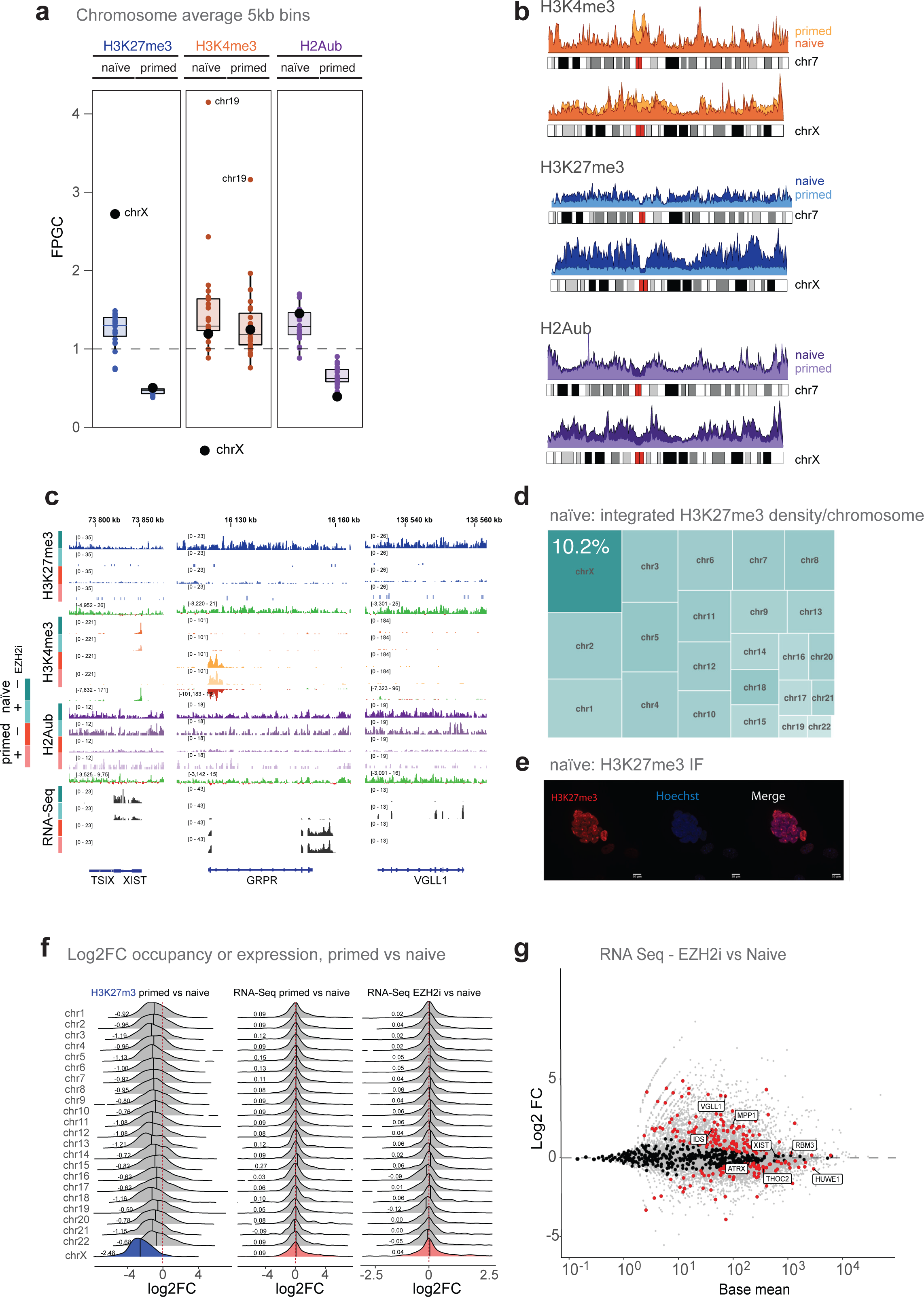
X chromosomes in naïve hESCs accumulate H3K27me3 without apparent chromosome-wide repressive function. **a** Chromosome average enrichment of H3K4me3, H3K27me3 and H2Aub in naïve and primed hESCs. RPGC of combined replicates are shown. **b** Chromosome density plot comparing the X chromosome signals in naïve and primed hESCs to those of an autosome with similar size (chr7). RPGC of combined replicates are shown. **c** Genome browser examples of regions on the X chromosome with differential occupancy of H3K27me3, H3K4me3 and/or H2Aub, with selected examples of genes expressed in naïve (XIST) or primed (GRPR) hESC. Signal shows three replicates combined, gain/loss tracks comparing naïve and primed signals. Tracks from the same histone modification are shown on the same RPGC scale. **d** Treemap showing a proportional representation of the total (integrated) amount of H3K27me3 by chromosome (area) and average density (color intensity). See also Extended Data Fig. 2b for additional controls. **e** Immunofluorescence microscopy showing nuclear H3K27me3 foci in naïve hESC. **f** Density plots of log2 fold-changes in promoter H3K27me3 levels (left) and RNA-Seq output (middle) of genes grouped by chromosome, comparing naïve and primed hESC. Density plots of log2 fold-changes in RNA-Seq output of genes grouped by chromosome, comparing untreated and EZH2i-treated naïve hESCs (right). Fold-changes were determined using DESeq2 from triplicate datasets. Median values by chromosome are given and indicated as vertical line in the density plot. **g** MA-plot of RNA-Seq data (Base mean and log2 fold-change as calculated with DESeq2 from triplicates) comparing untreated and EZH2i-treated naïve hESC. Genes on the X chromosome are highlighted in black (padj < 0.05 highlighted in red) with selected annotations.

We compared global levels of each histone modification across the four conditions as yielded by the input-normalized read counts ^33^ from MINUTE-ChIP. Global H3K4me3 levels were not significantly different between naïve and primed hESCs (Fig. 1b).

Notably, H3K27me3 levels were substantially (-3.3-fold) higher in naïve than in primed hESCs (Fig. 1b), an unexpected observation on the background of the existing body of literature reporting markedly low levels of H3K27me3 in the naïve state ^17, 27^. However our findings are consistent with more recent quantitative mass spectrometry data comparing naïve and primed hESCs ^26^. A higher H3K27me3 level in naïve hESCs was further confirmed by immunofluorescence microscopy (Extended Data Fig. 2b, c).

Global H3K27me3 levels were strongly depleted by treatment with EZH2i, with only 3.2% H3K27me3 signal remaining in naïve hESCs, and 7.6% in primed hESCs (Fig. 1b). Like H3K27me3, H2Aub signal was also significantly (2.1-fold) higher in naïve as compared to primed, suggesting a concerted regulation of PRC1/PRC2 activity. We sought to examine in more detail how the histone modification patterns distinguished naïve and primed hESCs. We performed genome-wide enrichment analysis on functionally annotated chromatin states (Fig. 1c, Extended Data Fig. 1b-d), as well as 10kb windows (Figures 1d). Despite no significant change in global levels (Fig. 1b, Extended Data Fig. 3a), H3K4me3 was increased somewhat at active promoters in the naïve state (Fig. 1c, Extended Data Fig. 3, 4). Bivalent promoters largely maintained stable H3K4me3 levels, but 11% showed a significant increase and 5% a decrease in H3K4me3 in the naïve state (Fig. 1c, Extended Data Fig. 3b, c).

**Fig. 3.**
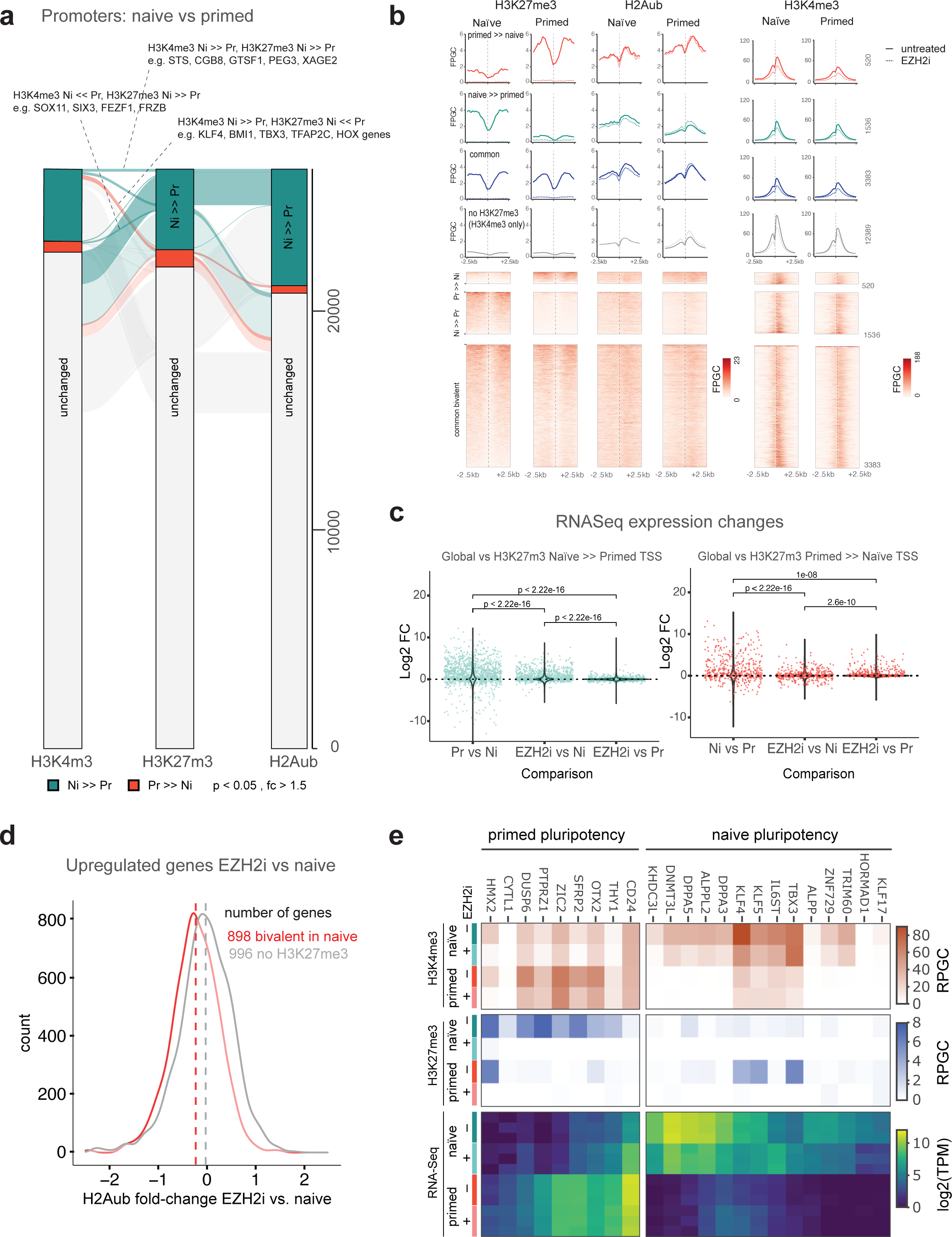
H3K27me3 is adaptive to gene expression changes between naïve and primed pluripotent states and contributes to repression of non-state specific genes. **a** Alluvial plot showing H3K27me3, H3K4me3 or H2Aub differentially occupied promoters (DESeq2 p.adj < 0.05 and fold-change > 0.5 from three replicates) between naïve and primed hESC, as well as the correspondence between the differentially occupied promoter groups across the three histone modifications. Functionally interesting connections (each containing more than 100 genes) are annotated.b *De novo* annotation of bivalent promoters based on DESeq2 analysis shown in A, B. Five promoter classes were defined: primed-only bivalent (Pr >> Ni), naïve-only bivalent (Ni >> Pr), common/shared bivalent, H3K4me3-only and H3K4me3 negative (not shown). Average H3K27me3, H2Aub, H3K4me3 profile plots in naïve and primed hESCs for each class and corresponding heatmaps for the first three states are shown. Additionally, profiles for EZH2i-treated naïve and primed conditions are shown as dashed lines. **c** Context-specific transcriptional response to global H3K27me3 depletion in different classes of bivalent genes. RNA-Seq changes (log2 fold-change from DESeq2 analysis from three replicates each) are plotted, comparing naïve and primed condition as well as EZH2i treatment and the respective control condition (left: naïve-only bivalent promoters, right: primed only bivalent promoter). The distribution of fold-changes of all genes is shown as violin plots, whereas the class-specific genes are shown as jitter points. Significance of pairwise Student’s t test is given. The mean fold-changes of the class-specific group was compared to the mean fold-change of all genes using Student’s t-test and Cohen’s d, and these values are given in the main text. See Extended Data Fig. 7B for corresponding analysis of the shared bivalent class of genes. **d** Density plot of fold-changes of H2Aub levels following H3K27me3 depletion in hESC. Only genes that were derepressed upon EZH2i-treatment (DESeq2 p.adj < 0.05, fold-change > 1.5 based on three replicates) were included in the analysis. The group of naïve-bivalent promoters (hence including promoters of the naïve-only and shared class) is compared to H3K27me3-devoid promoters. For an analysis of individual classes see Extended Data Fig. 7C. **e** Heatmap showing RNA-Seq expression levels (log2-transformed TPM) of previously defined marker genes for naïve or primed pluripotency ^64^ in naïve and primed hESCs (+/- EZH2i treatment), as well as the H3K4me3 and H3K27me3 levels (RPGC) at their respective promoter. RPGC from combined replicates are used for H3K4me3 and H3K27me3, whereas the three individual replicate TPM values are plotted for RNA-Seq data.

**Fig. 4.**
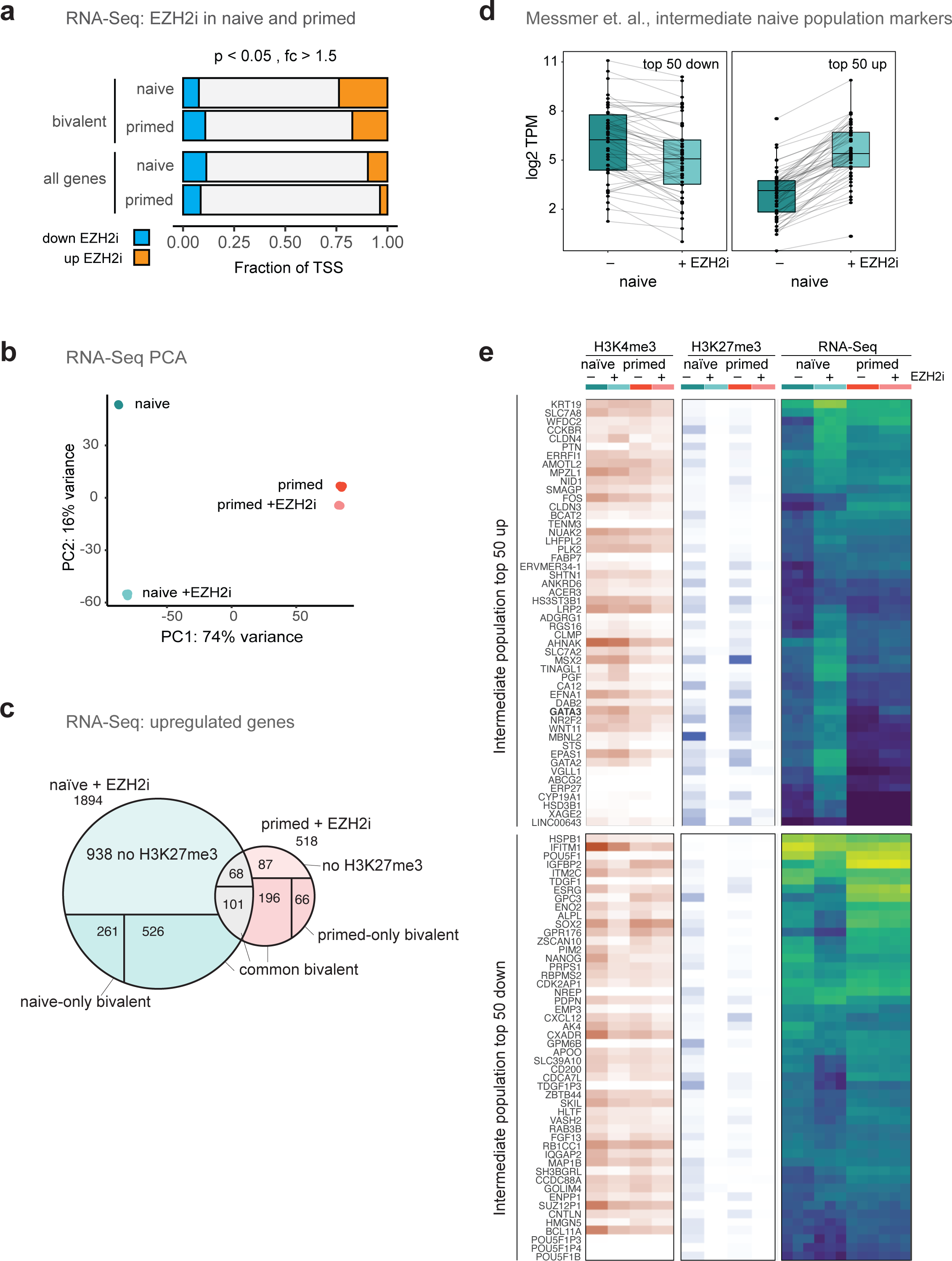
PRC2 inhibition derepresses a naïve-specific subset of bivalent genes. **a** Fractions of significant (p < 0.05) transcriptional changes in response to EZH2 inhibitor treatment of naïve and primed hESC, amongst all or annotated bivalent ^62^. **b** Intersection of genes derepressed after EZH2i treatment in naïve and primed hESC. Each venn intersection is further annotated by the H327me3 promoter class. **c** Principal component analysis of RNA-Seq datasets (triplicates shown individually). **d** Paired boxplots showing RNA-Seq expression changes (log2 TPM) of naïve cells in response to EZH2i treatment, for marker genes of an ’intermediate’ population present at -2% abundance in naïve cultures as defined in Messmer et. al. 2019 ^64^. The 50 top up- and down-regulated genes reported for the ’intermediate’ population were selected, and their expression in naïve hESCs with or without EZH2i treatment is shown. **e** Heatmap showing expression (TPM) in naïve and primed hESCs (+/- EZH2i treatment), as well as H3K4me3 and H3K27me3 promoter status of the same marker genes. RPGC from combined replicates are used for H3K4me3 and H3K27me3, whereas the three individual replicate TPM values are plotted for RNA-Seq data.

Naïve cells had a higher basal level of H3K27me3 across non-polycomb chromatin states and most of the 10kb bins (Figures 1c, d, Extended Data Fig. 3a). Broad H3K27me3 hypermethylation has been observed in mouse naïve ESC ^33, 35^ in the context of DNA hypomethylation ^35–38^. Global hypomethylation is also observed in human naïve pluripotency ^17, 22, 39^, and losses of CpG methylation coincided with gains in H3K27me3 across various chromatin states (Fig. 1c, Extended Data Fig. 2d), thus suggesting a similar interplay of H3K27me3 and DNA methylation in human ESCs. Despite the global gain, the average H3K27me3 level at bivalent promoters and polycomb-repressed chromatin states decreased mildly (Fig. 1c, Extended Data Fig. 1b). The average decrease was the result of a considerably changing H3K27me3 landscape at the level of individual genes (Fig. 1d, e, Extended Data Fig. 3): while more than 80% of promoters did not change H3K27me3 status (Extended Data Fig. 3b, c), approximately 5% of promoters including a number of *HOX* genes and other classical bivalent genes, showed reduced H3K27me3 in naïve hESCs (Fig. 1e, Extended Data Fig. 3b, c). Amongst these, naïve-specific genes such as *KLF2* and *KLF4* entirely lost H3K27me3 in the naïve state (Fig. 1e). On the other hand, more than 10% of promoters, e.g. *SOX11*, *DUSP6*, *OTX2* strongly gained H3K27me3 (Fig. 1d, e, Extended Data Fig. 3b, c). Our quantitative data thus contradicts earlier reports of a drastic loss of H3K27me3 across bivalent promoters in the naïve state ^17, 40, 41^.

### Broadly distributed H2A ubiquitination characterizes the naïve state

PRC1 is a powerful repressor complex essential for self-renewal, pluripotency and ordered differentiation of mESC ^42–45^. PRC1 acts largely independent of PRC2 in mESC ^46–49^. PRC1 subunit interaction maps and global binding patterns have been described in conventional hESCs and differentiated cell line models ^50–55^. However, the functional role of H2AUb in H3K27me3-dependent or independent transcriptional repression has not been studied in naïve or primed hESC.

Like H3K27me3, H2Aub showed a higher genome-wide occupancy in naïve hESCs (Fig. 1 c,d, Extended Data Fig. 3a-c), with less than 2% and -20% of promoters significantly lower and higher H2Aub levels in naïve, respectively (Extended Data Fig. 3b). While genome-wide H2Aub and H3K27me3-levels were reasonably correlated in primed and naïve hESCs, H2Aub did not mirror many of the gains and losses of H3K27me3 (Extended Data Fig. 3d). For example, levels of H2Aub did not significantly change between naïve and primed state for *DUSP6* and *OTX2* (Fig. 1e).

### Globally H2Aub landscape is largely independent of H3K27me3

To understand the functional role of PRC2 in setting up the naïve epigenomic and transcriptional landscape, we depleted H3K27me3 by inhibiting PRC2 in naïve and primed state. This allowed us to directly test whether H2A ubiquitination was dependent on H3K27me3 globally or locally. Intriguingly, global H2Aub levels were unchanged in H3K27me3-depleted naïve cells (Fig. 1b). Furthermore, the H2Aub landscape was maintained across all chromatin states, bivalent and active promoters (Fig. 1c), and less than 0.15 % of 10kb bins genome-wide showed a significant reduction in H2Aub, including -0.25% of annotated bivalent promoters (Extended Data Fig. 3e). For example, H2Aub levels were broadly reduced by 10-30% at *HOX* gene clusters (Fig. 1e), with maximal loss -50% at *HOXC5*, HOX*B8* and *HOXB9* promoters. Together, these results suggested that PRC1 recruitment and activity was largely independent of H3K27me3 with exception of a small set of bivalent genes, suggesting that this subset required H3K27me3 for efficient canonical PRC1 recruitment. In summary, our data suggested that H3K27me3 stands out as an adaptive mark shaping the unique epigenomes of naïve and primed pluripotent states.

### H3K27me3 accumulates on naïve X chromosomes

Surveying the chromosome-wide distribution of our quantitative ChIP-Seq profiles, we noted an overproportional H3K27me3 signal on the X chromosome of our naïve female H9 hESCs, which was also observed in other female hESCs (Extended Data Fig. 5a), transcriptional activity of both X-chromosomes, coupled to being coated by XIST as seen in the human pre-implantation embryos is considered one of the defining features of the naïve hESCs ^23, 56^. In cultured primed hESCs, a phenomenon termed X chromosome erosion (Xe) leads to a loss of XIST expression and H3K27me3 on the inactive X chromosome ^57^. The lack of enrichment for both H3K27me3 and H2Aub on the X chromosomes indicated an Xe state of ours and others’ cultured primed hESCs (Fig. 2a, b, Extended Data Fig. 5a).

**Fig. 5.**
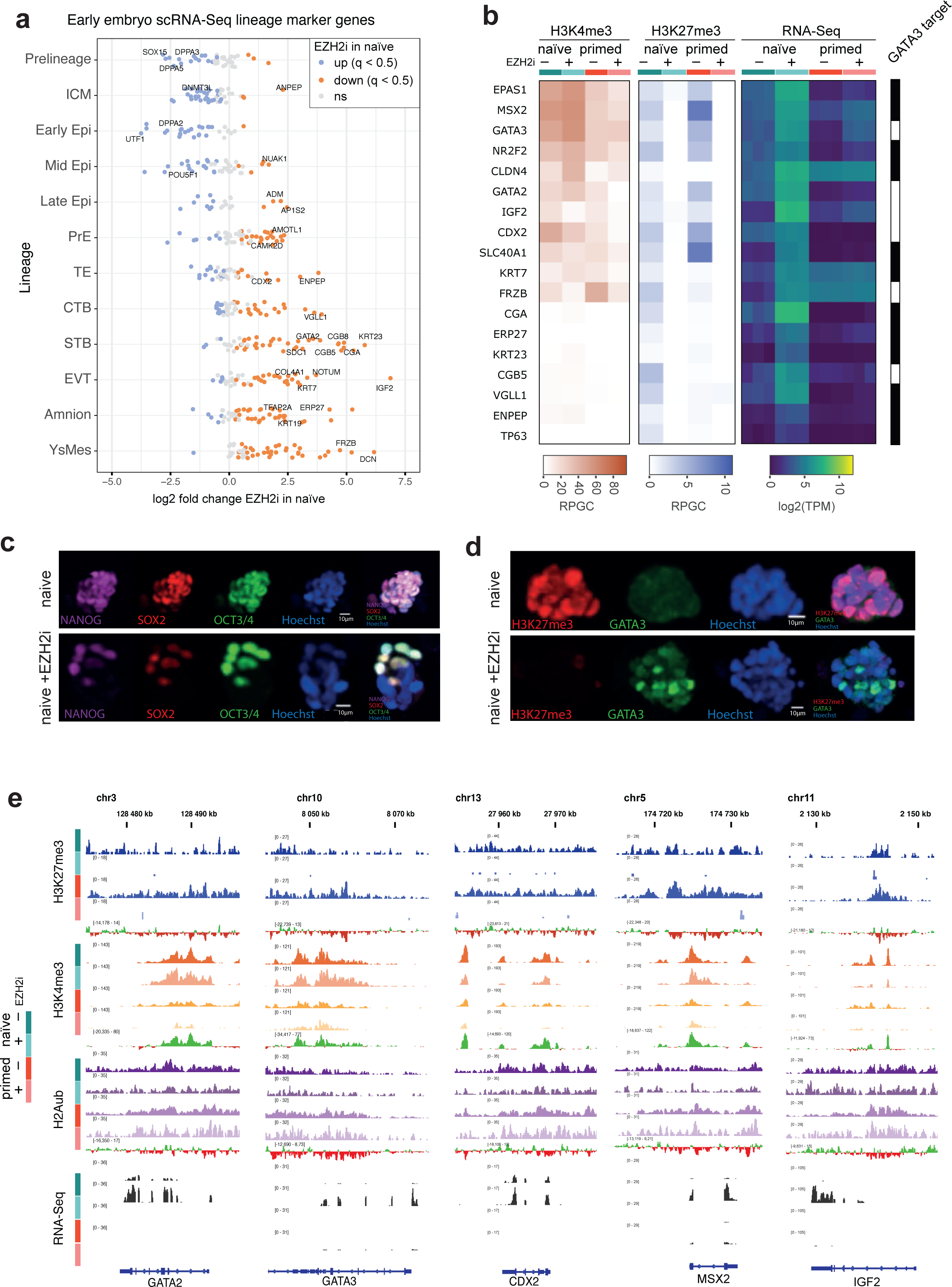
Loss of H3K27me3 in naïve activates trophectoderm and placental gene expression programs. **a** Stripchart showing expression of a comprehensive set of marker genes defined from human embryo single cell data comparing naïve hESCs and naïve hESCs treated with EZH2i. Markers are grouped into pre-lineage, inner cell mass (ICM), epiblast, primitive endoderm, (PrimEndo), trophectoderm (TE) cytotrophoblast (CTB), syncytiotrophoblast (STB), extravillous trophoblast (EVT), amnion and yolk sac mesoderm (YsMes) and primitive streak (PriS). Significant differences (DESeq2 p.adj < 0.05 from triplicate) are highlighted in color. **b** Heatmap showing expression (TPM) in naïve and primed hESCs (+/- EZH2i treatment), as well as H3K4me3 and H3K27me3 promoter status of selected trophectoderm and placenta-specific genes. RPGC from combined replicates are used for H3K4me3 and H3K27me3, whereas the three individual replicate TPM values are plotted for RNA-Seq data. GATA3 binding as determined by ChIP-Seq peaks during trophectoderm differentiation ^9^ is indicated. **c, d** Immunofluorescence microscopy of naïve hESC colonies with or without EZH2i treatment assessing relative expression of pluripotency marker (NANOG, SOX2, OCT3/4) as wells as trophectoderm transcription factor GATA3. **e** Genome browser examples of selected trophectoderm lineage markers. Shown is the combined signal of three replicates for H3K27me3, H3K4me3, H2Aub, RNA-Seq, as well as gain/loss tracks comparing naïve and primed signals. Tracks from the same histone modification are shown on the same RPGC scale. RNA-Seq expression is shown on the same TPM scale.

To further understand the nature and purpose of H3K27me3 enrichment on the active X chromosomes of naïve hESCs, we analyzed the distribution of H3K27me3, H3K4me3 and H2Aub on a chromosomal scale comparing naïve and primed states (Fig. 2a, b).

Strikingly, H3K27me3 density on X chromosomes was an outlier to an otherwise narrow distribution amongst the autosomes. H3K27me3 levels of the naïve X chromosomes were -2-fold higher than naïve autosomes and -5-fold higher than the average signal on primed chromosomes (Fig. 2a, b). At Mb scale, regions of very high H3K27me3 enrichment alternated with less enriched regions along the X chromosomes (Fig. 2b). In particular, we noted that transcriptionally active loci in the naïve state, such as the XIST locus, did not gain H3K27me3 as compared to the primed state (Fig. 2c, Extended Data Fig. 6a). Thus, H3K27me3 appeared to accumulate to high levels on the X chromosomes unless counteracted by transcription. Integrating the H3K27me3 signal across chromosomes, we estimated that the X chromosomes carry -10% of all H3K27me3 marks, more than any autosome (Fig. 2d), in concordance with immunofluorescence staining that showed distinctive H3K27me3 clouds for X chromosome (Fig. 2e) ^57^.

### H3K27me3 on naïve X chromosomes does not recruit H2Aub and does not contribute to dosage compensation

Notably, the exceptionally high levels of H3K27me3 on the naïve X chromosome were not mirrored by a similar chromosome-specific enrichment of H2Aub: the H2Aub signal from the naïve X chromosomes was not significantly different from autosomes (Fig. 2a, b). Moreover, depletion of H3K27me3 did not significantly change chromosome-wide H2Aub levels neither on autosomes nor the X chromosomes (Extended Data Fig. 5b). Thus, the exceptional accumulation of H3K27me3 on the naïve X chromosome does not appear to be targeted via H2A ubiquitination, and neither does high H3K27me3 levels cause an increase in H2Aub levels. This is in contrast to the role of H3K27me3 in X inactivation as studied in mESCs, where accumulation of H2Aub precedes and is required for accumulation of H3K27me3 ^58^.

Despite the strong increase in H3K27me3 comparing naïve and primed X chromosomes, RNA-Seq showed that overall transcriptional output from the X chromosomes relative to autosomes did not change, and the distribution of differentially expressed genes between naïve and primed states was similar to autosomes (Fig. 2f, Extended Data Fig. 6b). A phenomenon termed X chromosome dampening is thought to reduce dosage of the X chromosomes in the naïve state in a mechanism that is distinct from X inactivation and not well understood ^59^. Hence, we wondered if H3K27me3, in a non-canonical or H2Aub-independent manner, contributed to dosage compensation in naïve hESC, e.g. through chromosome-wide weak transcriptional repression that would attenuate output from the X chromosomes.

To follow up on this hypothesis, we analyzed chromosome-wide transcriptional output after EZH2i treatment of naïve and primed hESCs (Fig. 2f, g). Crucially, depletion of H3K27me3 using EZH2i in naïve hESCs did not lead to a general upregulation of transcription from the X chromosomes. Even though some genes, such as VGLL1 (Fig. 2c, g), were marked with H3K27me3 in the naïve state and strongly derepressed upon EZH2i treatment, up- and downregulated genes were similar in number and showed a similar fold-change range as those of autosomes (Fig. 2f, g, Extended Data Fig. 6c).

Thus, we conclude that H3K27me3 hypermethylation on the naïve X chromosome does not confer chromosome-scale dosage compensation. This does not exclude that H3K27me3, induced by XIST expression already on the naïve Xa, while functionally neutral to transcriptional output, may later promote X inactivation upon transition to primed state.

### Quantitative ChIP defines a new set of naïve-specific bivalent genes

Aside from the pronounced gain on the X chromosome, we noted above that a considerable number of gene promoters gained H3K27me3 in the naïve state (Fig. 1d,e, Extended Data Fig 3). However, the role of PRC2 in naïve hESCs has been elusive because prior studies reported overall low levels of H3K27me3 ^14, 16, 17, 40^ and that PRC2 activity was dispensable for naïve pluripotency ^60, 61^. Hence, we sought to revisit the role of PRC2 in repressing bivalent genes in naïve versus primed hESCs. Calling differential occupied promoters using DESeq2, 3712 promoters had significantly higher H3K27me3 levels in the naïve state, whereas 798 promoters had significantly higher levels in the primed state (Fig. 3a). Out of 4561 previously annotated bivalent genes in primed hESCs ^62^, 504 were significantly higher in naïve and 404 were higher in primed (Extended Data Fig. 7a).

Intersecting significant changes in H3K4me3 and H3K27me3 between naïve and primed hESCs indicated that they did not occur in a concerted manner (Fig. 1e, 3a), suggesting that H3K27me3 and H3K4me3 levels were controlled largely through independent mechanisms. However, three specific scenarios occurred at higher frequency (more than 100 promoters in each scenario) (Fig. 3a): the first set of genes were expressed only in the naïve state, such as *KLF4*, *KLF5*, *BMI1*, *TBX3*, *DNMT3L* lost H3K4me3 and gained H3K27me3 in the primed state. Another set of genes including *SOX11*, *SIX3* gained H3K4me4 and lost H3K27me3 in the primed state. A third set, including *STS*, *CGB8*, *XAGE2* had significantly higher levels of both H3K27me3 and H3K4me3 in the naïve state.

Since prior definition of bivalent gene sets were based on H3K27me3/H3K4me3 enrichment in primed hESCs only ^17, 62^, we called bivalent promoters *de novo* from our datasets in five categories based on their H3K27me3 enrichment: 520 primed-only,1536 naïve-only and 3383 common bivalent promoters, 12389 H3K4me3-only promoters, and a remaining large group of H3K4me3-negative promoters (not shown) (Fig. 3b, left).

H3K27me3 profiles reflected the expected primed-only, naïve-only or common enrichment. EZH2i treatment quantitatively removed H3K27me3 from all classes in naïve (>97.5% peak signal depletion) and primed hESCs (>95% peak signal depletion). H2Aub followed similar class-specific trends between naïve and primed states as H3K27me3 albeit within a smaller dynamic range (Fig. 3b, middle). Strikingly, as already hinted by the genome-wide analysis above (Fig. 1c, d), H2Aub was almost unperturbed in the absence of H3K27me3, showing the largest decrease (-15%) at bivalent promoters in the naïve state (Fig. 3b, middle).

H3K4me3 was co-enriched in both naïve and primed condition but tended to be higher in naïve state (Fig. 3b, right). Unexpectedly, H3K4me3 did not increase at any of the bivalent promoter classes upon EZH2i treatment (Fig. 3b, right). Instead H3K4me3 was reduced -2fold globally (Fig. 1b, Extended Data Fig. 4), and also at bivalent promoters (Fig. 3b, right). The EZH2 inhibitor used here, EPZ-6438, does not affect H3K4me3 levels in other cell types ^34^ and mESC (Extended Data Fig. 2c), thus suggesting that H3K27me3 depletion itself triggers a reduction of H3K4me3 specifically in hESC. We did not find evidence for a transcriptional downregulation of H3K4me3-specific histone methyl transferase complex (MLL) subunits or upregulation of corresponding demethylases that would provide a direct mechanistic explanation, hence the apparent link between H3K27me3 and H3K4me3 remains to be elucidated. We conclude that H3K27me3 is not generally antagonistic to H3K4me3. However, as detailed below, a specific subset of bivalent promoters indeed gained H3K4me3 upon H3K27me3 depletion, concomitant with a strong induction of transcription.

### H3K27me3 mediates repression of distinct sets of bivalent genes in naïve and primed hESC

Given the previously unappreciated naïve-specific set of bivalent promoters, we wondered if H3K27me3 played a functional role for these genes. To answer this question, we investigated our RNA-Seq data collected in the matching naïve and primed conditions, as well as EZH2i treatments.

We first focussed on the naïve-only and primed-only bivalent promoters where H3K27me3-mediated repression should be specific to the respective pluripotent state (Fig. 3c, d): naïve-only bivalent genes indeed were predominantly lower expressed (p < 5e-103, Cohen effect size medium) in naïve state than in primed (Fig. 3d, left panel), whereas primed-only bivalent genes were predominantly higher expressed (p = 3e-49, Cohen effect size large) in naïve (Fig. 3d, right panel, Extended Data Fig. 7b). Crucially, treating naïve hESCs with EZH2i derepressed naïve-only bivalent genes (p = 1e-77, Cohen effect size medium), while it did not significantly affect the expression of primed-only bivalent genes (Fig. 3d, left panel). Vice versa, treating primed hESCs with EZH2i derepressed primed-specific genes (p = 4e-75, Cohen effect size large) while not significantly affecting naïve-only bivalent genes (Fig. 3d, right panel). Finally, shared bivalent genes varied in expression level but as a group did not show a biased expression for either naïve or primed state (p = 1e-51, Cohen effect size small); EZH2i treatment increased expression in both states (Cohen effect size medium) (Extended Data Fig. 7c).

An important question that emanated from these observations was how H3K27me3 exerts the repression of bivalent genes, creating the poised state. The canonical pathway of Polycomb repression involves recruitment of canonical PRC1 complex via H3K27me3, following H2A ubiquitination, chromatin compaction and silencing. However H2Aub did not mirror forced depletion of H3K27me3, neither on a global level, nor at bivalent promoters (Fig. 1b, Fig. 3b). We thus asked if H2Aub would be affected more specifically at the subset of bivalent genes that were also transcriptionally derepressed upon H3K27me3 depletion: we observed a weak trend to lose a H2Aub (median loss 15%) at derepressed genes upon EZH2i treatment that were bivalent in the naïve state (i.e. from the naïve-only or common class) (Fig. 3f, Extended Data Fig. 7d). As a control, H2Aub did not follow this trend at H3K27me3-devoid promoters that were also upregulated with EZH2i (Fig. 3f, Extended Data Fig. 7d). Thus, a contribution of H2Aub to repression of bivalent genes in the naïve state is neither strongly supported nor refuted by our data.

We note that not all naïve-only and primed-only bivalent genes followed the trends described above (i.e. other naïve-only bivalent genes were not upregulated in primed state and/or with EZH2 inhibitor treatment). Hence, H3K27me3 must be considered as one part of a more complex regulatory network that ultimately determines transcriptional output. Nevertheless, our data reveals that promoter bivalency is not simply lost in the naïve state but redirected to a different set of genes. Promoter bivalency in the naïve state, as in the primed, has the capacity to create a transcriptionally poised state where H3K27me3 is required to maintain a repressed state.

### H3K27me3 contributes to basal repression of naïve and primed pluripotency markers

Amongst the genes switching bivalency status between naïve and primed state (Fig. 3b) were many known marker genes of naïve and primed pluripotency (see also examples above, *KLF4*, *OTX2*, *DUSP6* in Fig. 1e, Fig. 3a). Another key transcription factor uniquely expressed in naïve hESCs and recently implicated in the establishment of the naïve transcriptional landscape is TFAP2C ^63^. The promoters of *TFAP2C* gene, as well as the related *TFAP2A* gene were indeed highly enriched in H3K4me3 and devoid of H3K27me3 in the naïve state, but acquired a bivalent state with intermediate H3K4me3 levels and high H3K27me3 enrichment in the primed state (Extended Data Fig. 8a).

Intriguingly, depleting H3K27me3 in the primed state basally activated *TFAP2A/C* transcription, albeit not to the same high level observed in the naïve. Thus, depletion of H3K27me3 in primed cells removed PRC2-mediated repression but was not able to revert the *TFAP2A/C* genes to a full ’on’ state, presumably because additional activating factors were not expressed in the primed state.

To generalize this observation, we assessed H3K27me3 and H3K4me3 status, as well as RNA-Seq output for comprehensive sets of naïve and primed markers recently derived from scRNA-Seq ^64^. We observed a similar switch from H3K4me3-only promoter status in naïve to bivalent in primed for three additional transcription factors implicated in setting up the naïve pluripotency (*KLF4*, *KLF5*, *TBX3*), and these also increased basal transcription in response to H3K27me3 depletion in the primed state (Fig. 3e). However, the majority of naïve markers, such as *DNMT3L*, *TRIM60*, *ZNF729*, did not accumulate H3K27me3 in the primed state and were also not responsive to EZH2i treatment (Fig. 3e).

On the other hand, most markers of primed pluripotency ^64^ had H3K4me3-only promoters in the primed state, while assuming a H3K27me3-high/H3K4me3-low bivalent promoter status in the naïve state (Fig. 3e). H3K27me3 depletion in the naïve state resulted in increased transcription of most of these genes (including *OTX2*, *CD24*, *DUSP6*), albeit in no case reaching the levels of the primed state (Fig. 3e, see also Fig. 1e). Hence, we identify a new function for PRC2 in naïve pluripotency, repressing basal expression of primed-specific genes.

Concomitant with induction of primed-specific factors, naTve markers including *DPPA3*, *KLF4*, *KLF5*, *DNMT3L* (Fig. 3e), as well as core pluripotency factors (Extended Data Fig. 8c) were downregulated. In conclusion, H3K27me3 contributes to repression of primed-specific factors in the naïve state, and to a limited extent also to the repression of naïve-specific factors in the primed state (Fig. 3e). Hence, PRC2 appears to establish an epigenetic barrier between the two states through adaptive repression of different gene sets in naïve and primed pluripotency. Viewed in a developmental trajectory, this may suggest that genes important for primed pluripotency are maintained in a poised state during naïve pluripotency, to be activated rapidly at implantation.

### PRC2 inhibition derepresses a naïve-specific subset of bivalent genes

Having so far focussed on genes that switch bivalent state between naïve and primed, we considered that a wider set of bivalent promoters may be significantly repressed by H3K27me3 in the naïve state, and examined all differential expressed genes between EZH2i treated and untreated naïve state. Approximately similar numbers of genes were up- and downregulated upon EZH2i treatment, and, as already noted above, bivalent genes were predominantly upregulated (Fig. 4a). Principal Component Analysis (PCA) of our RNA-Seq datasets showed that EZH2i treatment had a much more profound effect in naïve cells (Fig. 4b). Out of 1894 upregulated genes (log fold-change > 1.5, FDR < 5%), 898 were enriched in H3K27me3 in their promoter region in the naïve state (Fig. 4c). 261 of these were previously categorized as naïve-only H3K27me3 promoters (Fig. 3b), hence we considered these genes bona-fide naïve-specific targets of PRC2, as already discussed above. On the other hand, 349 genes were uniquely upregulated with EZH2 inhibitors in the primed state, and 262 out of these were H3K27me3-enriched (Fig. 4c). An additional 169 genes were upregulated in both states and 101 of these also were bivalent in both states (Fig. 4c). The small proportion of co-regulated genes amongst the genes featuring bivalent promoters in both states highlights the exquisitely context-specific function of PRC2 in repressing largely non-overlapping gene sets in closely related pluripotent states (Fig. 4c). Evidently, additional context-dependent cues must determine if loss of H3K27me3 at bivalent promoters is sufficient to switch the respective gene on.

### H3K27me3-depleted naïve hESCs show similarities to a rare subpopulation observed in normal naïve hESCs cultures

Beyond the group of H3K27me3-enriched genes that may represent direct targets of PRC2 repression, we observed an additional 938 upregulated and a similar number of downregulated genes without notable H3K27me3 enrichment (Fig. 4c). This suggested a wider transcriptional reprogramming triggered by depletion of H3K27me3, and we explored possible correlations with known embryonic expression signatures. A recent single-cell RNA-Seq study profiled naïve and primed hESCs cultures ^64^ and identified a subpopulation amongst naïve hESCs that gained expression of some markers of primed hESCs, as well as a number of unique genes absent in either naïve or primed state, including *ABCG2*, *CLDN4*, *VGLL1*, *GATA2*, *GATA3*, and *ERP27* . We initially noted that the marker genes for this so-called ’intermediate’ population were also expressed in our EZH2i-treated naïve cells (e.g. *VGLL1*, Fig. 2g). Strikingly, all top 50 upregulated genes reported for the intermediate population were also amongst the top-most upregulated genes in our EZH2i-treated naïve cells, and most of the top 50 downregulated genes were also downregulated (Fig. 4d). An analysis of promoter status revealed that many of the upregulated marker genes were bivalent in the naïve state, transitioning to a H3K4me3-only state with EHZ2i treatment (Fig. 4e, Extended Data Fig. 9). Hence, our data suggested that EZH2 inhibition can drive naïve cells towards a transcription program that is also naturally exhibited by a small subpopulation within cultured naïve hESCs. Representing only -2% of cells in a naïve hESC culture ^64^, the identity of these cells has however not been explored.

### PRC2 is required for repression of trophectoderm and extraembryonic tissue markers in naïve hESC

To functionally define the gene expression signature we identified, we further mined a comprehensive list of lineage markers derived from scRNA-Seq of the developing human embryo ^65^. A striking pattern emerged assessing the expression of these marker genes in EZH2i-treated naïve cells: Inner cell mass, blastocyst and epiblast markers were consistently downregulated while trophectoderm, extraembryonic and placental markers - cytotrophoblast (CTB), syncytiotrophoblast (STB), extravillous trophoblast (EVT), amnion and yolk sac mesoderm - were upregulated (Fig. 5a, e). The induction of these marker genes upon EZH2i treatment was specific to the naïve state (Fig. 5b, Extended Data Fig. 10a). Amongst the most highly upregulated marker genes was *ENPEP*, the gene for a surface marker (APA) that marks trophectoderm progenitor cells capable of forming STB ^9, 66^; *ABCG2*, a plasma membrane transporter highly expressed in trophectoderm and human placenta ^5, 67^; *TP63*, a p53 family member defining the CTB stem cell compartment ^5, 68^. Well known transcription factors specifying trophectoderm lineage, CDX2 ^69, 70^, GATA3 ^9, 70, 71^, MSX2 ^72^, NR2F2 ^73^ were also specifically upregulated in EZH2i treated naïve hESCs (Fig. 5b-e).

GATA3 has been considered the master transcription factor for trophectoderm development. GATA3 overexpression in hESCs can replace BMP4 stimulation to induce trophectoderm development ^9, 71^. GATA3 has also been shown to repress pluripotency transcription factors ^9^. Further, ChIP-Seq in trophoblast progenitors has shown that GATA3 binds to the promoters of many marker genes of trophectoderm and placental development, including *KRT23*, *VGLL1*, *CGA* and *TP63* ^9^. Intersecting with the published binding data, we found 20% of upregulated genes in EZH2i treated naïve cells to be bound by GATA3 (Extended Data Fig. 10b).

Investigating the promoter status of individual induced genes, we noticed two distinct groups - one group of promoters was H3K4me3/H3K27me3 bivalent, one had no H3K4me3 and typically also low or no H3K27me3 (Fig. 5b, e, Extended Data Fig. 10b). It has been proposed that GATA3 and other bivalent TF genes are at the top of a hierarchy of gene regulatory events leading to an induction of trophectoderm development ^9^. Indeed, those marker genes that did not carry H3K4me3 in the naïve state often featured GATA3 binding sites, suggesting they may be directly activated by GATA3 (Fig. 5b).

We sought to confirm GATA3 induction on the protein level and stained naïve and EZH2i-treated naïve hESC colonies with NANOG, SOX2, OCT4 and GATA3. Notably, EZH2 inhibition induced heterogeneity within colonies, with individual cells losing all three pluripotency markers while others maintained normal levels (Fig. 5c). This was concomitant with the appearance of GATA3+ single cells (Fig. 5d). Primed hESCs stained and imaged alongside the naïve samples did not show a loss of pluripotency markers or GATA3 expression with EZH2 inhibition (Extended Data Fig. 10c).

The induction of GATA3 positive and NANOG/SOX2/OCT4 negative cells suggests that in the course of the 7 day EZH2i treatment, a fraction of naïve hESCs have committed to trophectoderm lineage and exited from pluripotency. A time course experiments of BMP4-induced trophectoderm development from primed hESCs ^9^ established a temporal sequence of marker genes, with *GATA3* and *MSX2* being induced already after 8 hours, while *TP63*, *VGLL1* and *CGA* are only induced after 48 and 72h respectively (Extended Data Fig. 10d) ^9^. The expression of *TP63*, *VGLL1*, *CGA* in our 7d EZH2i treatment hence suggests that, once committed, the GATA3-positive cells are able to execute a typical trophectoderm development program towards late trophectoderm/STB markers (Extended Data Fig. 10d, 11).

In summary, chromatin and transcriptomic signatures suggest that depleting H3K27me3 drives a fraction of naïve hESCs into spontaneous commitment to a trophectoderm fate. Hence, PRC2 adopts an important role in shielding naïve pluripotent cells from aberrant trophectoderm differentiation.

## DISCUSSION

Our findings disagree in some aspects with earlier reports that PRC2 is dispensable for maintenance of naïve pluripotency in 5i/L/A and t2iGo media ^60, 61^. On the one hand, we find that EZH2i-treated naïve hESCs maintain typical domed colony growth thus suggesting that trophectoderm commitment is a stochastic event that results in only a fraction of cells having exited pluripotency at any given time. In support, we observe most cells within a colony maintaining pluripotency marker expression at normal levels, while a subset loses pluripotency and gains GATA3 expression (Fig. 5c, d). On the other hand, we observe an overall reduction of core pluripotency marker expression on the population level (Extended Data Fig. 7d), while Shan et al. ^61^ have reported that NANOG, OCT4, SOX2 are unaffected in naïve EZH2-/- hESCs with additional shRNA-mediated depletion of EZH1. However, in this study, remaining EZH1 or H3K27me3 levels were not explicitly assessed, leaving the possibility that our EZH2i treatment resulted in a greater depletion of H3K27me3, thus a more pronounced phenotype. Of note the EZH2 inhibitor EPZ-6438 also inhibits EZH1 with nanomolar IC50 ^74^, likely achieving complete inhibition of EZH1 and EZH2 at the 10µM concentration used here (Fig. 1b, 3b).

Capturing quantitative differences between epigenomic profiles, as achieved with MINUTE-ChIP, has allowed us to uncover a more complex pattern of gains and losses of H3K27me3 between naïve and primed states than previously appreciated ^14, 16, 17, 40^. Importantly, we found that the bivalency was largely maintained in the naïve state and found hundreds of promoters that gained bivalency uniquely in the naïve state. Surprisingly, despite the extensive overlap of target promoters between naïve and primed states, inhibiting PRC2 catalytic activity activated independent gene sets, characteristic for each state. Hence, H3K27me3 per se cannot be attributed a constitutive repressive function at bivalent promoters, instead we view bivalency as a default state of many developmental genes that can optionally acquire a transcriptionally poised state (i.e. an state of low transcription in which solely H3K27me3 impedes activation) in the presence of cell type/lineage specific factors.

Such context-specific poising can explain why naïve but not primed hESCs cells gain trophectoderm differentiation potential when treated with EZH2i. Primed hESCs can give rise to trophectoderm ^9, 66, 68, 71^, demonstrating the capacity to transdifferentiate. However, genetic ablation of PRC2 in primed hESCs results in meso-endoderm development rather than trophectoderm development ^31, 61^. It is important to note that the process of exiting pluripotency and meso-endoderm speciation is not a rapid response to depletion of PRC2 in primed hESC, but emerges over 3-4 weeks of culture ^31, 61^. Over the course of our 7d EZH2i treatment we do not observe a loss of core pluripotency markers or upregulation of endo-mesoderm markers such as GATA6, GATA4 ^75^ (Extended Data Fig. 8b, 10a), in line with a time course conditional PRC2 knockout experiment showing first induction of endo-mesoderm at day 12 and decrease in pluripotency markers at day 20 ^61^. Therefore, the transcriptional rewiring and appearance of GATA3-positive/pluripotency-negative cells within 7 days of EZH2i treatment of naïve hESCs is in striking contrast to the slow exit from pluripotency of primed hESCs. Together with the observations that a subpopulation with expression of trophectoderm markers is already present in naïve cultures ^22, 64^ and that trophectoderm can be derived directly from naïve hESCs ^6, 76–78^, our data supports that the naïve state is uniquely poised towards trophectoderm lineage. We find that key trophectoderm transcription factors and other marker genes upregulated upon loss of H3K27me3, including *CDX2*, *GATA3*, *EPAS1*, *MSX2*, *TBX3*, *STS*, *CGA*, *CGB8* are already basally expressed in naïve hESCs while absent in primed (Fig. 3e, Fig. 5b, Extended Data Fig. 11). The bivalent promoters of these genes feature higher H3K4me3 and lower H3K27me3 levels in the naïve state as compared to primed, suggesting that activating signals present uniquely in the naïve state skew the poised state towards activation, while H3K27me3 acquires a crucial role in opposing full activation.

A candidate transcription factor for setting up the tight interconnectivity between naïve pluripotency and trophectoderm in humans is TFAP2C. While instrumental for setting up the transcriptional landscape of human naïve hESCs ^63^, TFAP2C is also a well-studied driver of trophectoderm development in mouse and is highly expressed throughout extraembryonic tissues including human and mouse placenta ^5, 9, 79^. TFAP2C is one of the key transcription factors induced when converting primed hESCs to trophectoderm by BMP4 stimulation, binding to promoters or enhancers of many trophectoderm genes ^5, 9^. TFAP2C is expressed -8fold higher in naïve hESCs as compared to primed and its expression does not further increase upon EZH2i treatment (Extended Data Fig. 8a). Hence, naïve cells already express a critical amount of TFAP2C to facilitate trophectoderm differentiation. Many trophectoderm marker genes are bivalent throughout naïve and primed pluripotency. The absence of activating transcription factors like TFAP2C, GATA2, GATA3 in the primed state might ultimately explain why H3K27me3 depletion does not result in increased expression of these PRC2 target genes. Only in the presence of a critical mass of trophectoderm transcription factors, PRC2 activity might become crucial in setting up an epigenetic barrier to prevent spontaneous commitment to trophectoderm differentiation. We speculate that even in standard naïve cultures, this epigenetic barrier is crossed in single cells with very low frequency. Once overcome, the existing pool of transcription factor proteins can rapidly gain control of the transcriptional program, antagonising the core pluripotency network.

In summary, our quantitative comparison of epigenetic landscapes in naïve and primed hESCs reveals an extensive rewiring of promoter bivalency in which PRC2-mediated H3K27me3 constitutes an important epigenetic barrier that dynamically follows and reinforces lineage choices and developmental progression. It will be an exciting focus of future work to elucidate if and how PRC2 is involved in guiding the totipotency-to-pluripotency transition in the early human embryo and if it can be utilized to facilitate establishment of human blastoids.

## MATERIALS AND METHODS

### Data and Software availability

The high-throughput data reported in this study have been deposited in GEO under the accession number <u>GSE181244</u>, which includes demultiplexed and deduplicated reads and a quantitatively scaled bigwig track for each sample. Additional code, supplementary data and html summaries are available on GitHub: https://github.com/elsasserlab/hesc-epigenomics

### Culture of human embryonic stem cells

Primed H9 (Wicell; WA09) p28(11) were thawed using Nutristem hESC XF medium (Biological Industries; 05-100-1A) and were cultured on tissue culture plates (Sarstedt; 83.3922) pre-coated with 10% LN521 (Biolamina; LN521-02). Passaging was done when confluency reached 70-80%. Naïve H9 p28(8) naïve p7 that had been previously converted to the naïve stem cell state using NaïveCult^TM^ induction media (Stemcell Technologies; 05580) were thawed onto high-density mouse (ICR) inactivated embryonic fibroblasts (MEFs, Gibco; A24903) plates using NaïveCult expansion media (Stemcell Technologies; 05590) supplemented with 10µM ROCKi (Merck; Y-27632). Twenty-four hours after thawing media was changed to fresh NaïveCult expansion media. Passaging was done every four to five days with media supplemented with 10µM ROCKi. Both hESCs were cultured at 37°C, 5% O_2_ and 5% CO_2_. For the EZH2i treatment, primed and naïve cells were grown in respective media, supplemented with 10µM EZSolution™ EPZ-6438 (BioVision; 2428-5) for seven days, with daily media changes. For sample collection, primed and naïve H9 were dissociated using TrypLE Select (Gibco; 12563011). Primed cells were washed once with phosphate buffered saline (PBS; Sigma D8537) and counted using Moxi Z mini automated cell counter (Orflo; MXZ001). Harvested naïve cells were centrifuged at 300 x g for 4 min, resuspended in fresh NaïveCult expansion media and kept on ice. Mouse feeder removal microbeads (Milteny Biotech; 130-095-531) were used according to the manufacturer’s protocol to reduce the amount of MEFs in the naïve cell samples. In brief, naïve cells were mixed with the microbeads and incubated at 4°C for 15 min. Meanwhile, the columns were equilibrated with NaïveCult media. The cell-microbead suspension was added to the column mounted onto a magnet, allowing unbound naïve cells to pass through into a collection tube. Columns were rinsed using NaïveCult media, and the flow-through was collected into the same collection tube. Counting was performed using the Countess™ II Automated Cell Counter (Applied Biosystems; A27977).

### Chromatin Immunoprecipitation (ChIP) sequencing

Triplicate pellets of 1x10^6^ cells were collected for all conditions, flash frozen and stored at -80°C prior to use. Mouse feeder removal microbeads (Milteny Biotech; 130-095-531) were used according to the manufacturer’s protocol to reduce the amount of MEFs in the naïve cell samples. Samples were prepared for ChIP-seq following the MINUTE ChIP protocol ^33^. Briefly native cell pellets were lysed, MNase digested to mono- to tri- nucleosome fragments and ligated with dsDNA adaptors (containing T7 promoter, 8bp sample barcode and a 6bp unique molecular identifier (UMI)) in a one pot reaction. Barcoded samples were then pooled and aliquoted into individual ChIP reactions with Protein A/G magnetic beads (BioRad; 161- 4013/23) coupled with the desired antibodies (5 ug each of H3K27me3 {Millipore 07-449}, H3K4me3 {Millipore 04-745} and H2AUb {Cell Signaling 8240S}) . Upon incubation for 4 h at 4°C with rotation, ChIP DNA was isolated and set up in sequential reactions of *in vitro* transcription, RNA 3’ adapter ligation, reverse transcription and PCR amplification to generate final libraries for each ChIP (Extended Data Fig. 1a). After quality assessment and concentration estimation, libraries were diluted to 4nM, combined and sequenced on the Illumina NextSeq500 platform with paired-end settings.

### RNA sequencing

1x10^6^ cells per growth condition were harvested, resuspended in Buffer RLT (Qiagen; 74106), spiked in with 5x10^4^ Drosophila cells per sample and then total RNA was extracted using the RNeasy Plus Mini Kit (Qiagen; 74136) according to the manufacturer’s protocol. Mouse feeder removal microbeads (Milteny Biotech; 130-095-531) were used according to the manufacturer’s protocol to reduce the amount of MEFs in the naïve cell samples. Purified RNA quantities were estimated using the Qubit RNA HS assay kit (Life technologies; Q32852) and the samples were subsequently flash frozen. RNA-seq libraries were generated and sequenced through BGI service (www.bgi.com) for strand specific RNA-seq with PolyA selection (DNBseq Eukaryotic Transcriptome De novo Sequencing).

### Immunofluorescence

Four percent PFA-fixed samples were permeabilized using 0.3% Triton X-100 (Sigma Aaldrich; T9284-100ML) in PBS (Gibco; 14190144) for 10 min, after which three washes with PBS were carried out. The samples were blocked for two hours using 0.1% Tween-20 (Sigma Aaldrich; P9416-100ML) and 4% FBS (ThermoFisher; 10082147) in PBS. Primary antibodies (GATA3 clone L50-823 (1:200, BD; 558686), H3K27me3 C36B11 (1:500, Cell Signaling Technologies; 9733S), OCT4 (1:200, SantaCruz; sc-5279), SOX2 clone EP103 (1:3, Biogenex; AN833) and NANOG (1:200, RnD; AF1997-SP)) were diluted in blocking solution and added to the samples, which were incubated at 4°C overnight. Excess antibodies were washed away using blocking buffer. Secondary antibodies:, donkey a-mouse IgG (H+L) Alexa fluor 555, donkey a-rabbit IgG (H+L) Alexa fluor 647, donkey a-goat IgG (H+L) Alexa fluor 647, donkey a-mouse IgG (H+L) Alexa fluor 488, donkey a-rabbit IgG (H+L) Alexa fluor 555 (all from Thermofisher; A-31570, A-31573, A-21447, A-21202 and A-31572, respectively) were diluted in blocking solution, added to the samples and then incubated for two hours in RT. Again, excess antibodies were washed away and samples were incubated with Hoechst 33342 (Thermofisher; H3570), which was followed by another set of washes. Samples were mounted using DAKO fluorescent mounting media (DAKO; S3023). Images were acquired using a Nikon Eclipse Ti spinning disk confocal microscope with a 20X air and a 60X oil immersion objective, respectively, and Z-stacks were analyzed using ImageJ.

### Immunoblotting

1x106 cell pellets for each growth condition were lysed in 100 µl of ice-cold radioimmunoprecipitation (RIPA) buffer (0.1% sodium deoxycholate, 0.1% SDS, 1% Triton X-100, 10 mM Hepes [pH 7.6], 1 mM EDTA, 5% glycerol and 140 mM NaCl) supplemented with Protease Inhibitor Cocktail (PIC, Roche) on ice for 10 min. Lysates were homogenized by sonication for 8-10 cycles at high power, 30 seconds on/off in a Bioruptor sonicator (Cosmo Bio Co. Ltd.). Samples were boiled at 95 °C for 5 mins with 6xSDS sample buffer before loading onto 4-20% Tris-glycine gels (BioRad). Resolved proteins were transferred to nitrocellulose membranes using the Trans-Blot® Turbo system (BioRad) according to the manufacturer’s instructions. Membranes were then blocked for 1 h in 1% casein prepared in Tris-buffered saline and 0.1% Tween-20 (TBS-T) before blotting with respective primary antibodies diluted in TBST, overnight at 4°C. Blots were washed three times with TBST and incubated with secondary antibodies in the same buffer for 1 h at room temperature (protect from light). After three TBST washes, the membranes were imaged on a LI-COR Odyssey ® FC system. Quantitation of signal and analysis was performed using the LI-COR Image studio software. Primary antibodies included total H3 1:10,000 (Active motif 39763), H3K4me3 1: 5000 (Millipore 04-745), H3K27me3 1: 5000 (Millipore 07-449) and H2Aub (Cell Signaling 8240S). The secondary antibodies were IRDye® 680RD anti-rabbit and IRDye® 800CW anti-mouse (LI-COR) at 1:5000 dilution.

### MINUTE-ChIP analysis

Preparation of FASTQ files

Sequencing was performed using 50:8:34 cycles (Read1:Index1:Read2) Illumina bcl2fastq was used to demultiplex paired-end sequencing reads by 8nt index1 read (PCR barcode). NextSeq lanes were merged into single fastq files, creating the primary fastq files. Read1 starts with 6nt UMI and 8nt barcode in the format NNNNNNABCDEFGH.

### Primary analysis

MINUTE-ChIP multiplexed FASTQ files were processed using minute, a data processing pipeline implemented in Snakemake ^80^. In order to ensure reproducibility, a conda environment was set. Source code and configuration are available on GitHub: https://github.com/NBISweden/minute. Main steps performed are described below.

### Adaptor removal

Read pairs matching parts of the adaptor sequence (SBS3 or T7 promoter) in either read1 or read2 were removed using cutadapt v3.2 ^81^.

### Demultiplexing and deduplication

Reads were demultiplexed using cutadapt v3.2 allowing only one mismatch per barcode. Demultiplexed reads were written into sample-specific FASTQ files used for subsequent mapping and GEO submission.

### Mapping

Sample-specific paired FASTQ files were mapped to the human genome (hg38) using bowtie2 (v2.3.5.1) with --fast parameter. Alignments were processed into sorted BAM files with samtools (v1.10). Pooled BAM files were generated from replicates using samtools.

### Deduplication

Duplicate reads are marked using UMI-sensitive deduplication tool je-suite (v2.0.RC) (https://github.com/gbcs-embl/Je/). Read pairs are marked as duplicates if their read1 (first-in-pair) sequences have the same UMI (allowing for 1 mismatch) and map to the same location in the genome. Blacklisted regions were then removed from BAM files using BEDTools (v2.29.2).

### Generation of coverage tracks and quantitative scaling

Input coverage tracks with 1bp resolution in BigWig format were generated from BAM files using deepTools (v3.5.0) bamCoverage and scaled to a reads-per-genome- coverage of one (1xRPGC, also referred to as ’1x normalization’) using hg38 genome size 3095978588. ChIP coverage tracks were generated from BAM files using deepTools (v3.5.0) bamCoverage. Quantitative scaling of the ChIP-Seq tracks amongst conditions within each pool was based on their Input-Normalized Mapped Read Count (INRC). INRC was calculated by dividing the number of unique hg38-mapped reads by the respective number of Input reads: #mapped[ChIP] / #mapped[Input] . This essentially corrected for an uneven representation of barcodes in the Input and we previously demonstrated that the INRC is proportional to the amount of epitope present in each condition ^33^. Untreated naïve hESCs (pooling replicates) were chosen as the reference condition, which was scaled to 1x coverage (also termed Reads per Genome Coverage, RPGC). All other conditions were scaled relative to the reference using the ratio of INRCs multiplied by the scaling factor determined for 1x normalization of the reference: ( #mapped[ChIP] / #mapped[Input] ) / ( #mapped[ChIP_Reference] / #mapped[Input_Reference] ) * scaling factor.

### Quality control

FastQC was run on all FASTQ files to assess general sequencing quality. Picard (v2.24.1) was used to determine insert size distribution, duplication rate, estimated library size. Mapping stats were generated from BAM files using samtools (v1.10) idxstats and flagstat commands. Final reports with all the statistics generated throughout the pipeline execution are gathered with MultiQC ^82^.

### Downstream analysis and visualization

Total mapped read counts from BAM files were used to calculate relative global levels of histone modifications. Summary values for fixed sized bins or custom intervals were calculated from scaled BigWig files using wigglescout (https://github.com/cnluzon/wigglescout). Differentially H3K27m3-enriched bins and gene TSS were calculated using DESeq2 (v1.32.0) using fixed size factors since bigWig files were already scaled by the minute pipeline. Figures were created using R (v4.1.0) ggplot2 (v3.3.5). Additionally, combined heatmaps were created using heatmaply (v1.2.1) and extra statistics were plotted with ggpubr (v0.4.0) package when required. Fig. 2d was made with ggridges (v 0.5.3). Fig. 2b was made with karyoploteR package (v1.18.0) ^83^. Fig 3a was done with ggalluvial (v 0.12.3). Genome track figures were made using IGV image export function. Corresponding source code for data analysis and figures can be found in the GitHub repository companion to this publication (https://github.com/elsasserlab/hesc-epigenomics). The repository was built using workflowr (v1.6.2). Data analyses were rendered from R markdown notebooks and results can be navigated at the corresponding website (LINK), generated using workflowr package (v1.6.2) ^84^.

### RNA-Seq analysis

#### Primary analysis

RNA-seq data was analysed by RNA-seq pipeline (v2.0) available in nf-core (https://nf-co.re/rnaseq/2.0) with hg38 as reference, using STAR ^85^ as read aligner and RSEM ^86^ to quantify read counts.

#### Downstream analysis and visualization

Read counts produced by RSEM were used as input for DESeq2 ^87^ differential expression analysis with default parameters. Log2FC shrinkage apegm ^88^ was used to filter low read count genes. Significance adjusted p-value cutoff was set on p < 0.05 and a fold change of 2. In heatmap RNA-seq figures, Transcripts Per Million (TPM) values are shown as (log2(TPM) + 1) instead of raw counts. Figures were created using R (v4.1.0) ggplot2 (v3.3.5). Additionally, combined heatmaps were created using heatmaply (v1.2.1) and extra statistics were plotted with ggpubr (v0.4.0) package when required.

#### Marker-gene detection for lineages

Published datasets from E-MTAB-3929^23^, GSE136447 ^89^, and http://www.human-gastrula.net/ ^90^were downloaded and processed as described previously^65^. Briefly, raw reads were mapped to hg38 reference genome using the STAR aligner with default settings, and only uniquely mapped reads were kept for gene expression quantification. Read counts were further estimated using rsem-calculate-expression from RSEM tool with the option of “--single-cell-prior”. Rescaled log-normalized counts using the deconvolution strategy implemented by the computeSumFactors function in R scran package^91^ and followed by multiBatchNorm function in the R batchelor package^92^ were used to detect the marker genes. Cells belong to hemogenic endothelial progenitors and erythroblasts from Carnegie stage 7 were excluded from the analysis. Also, cells labeled as “ICM” and “PSA-EPI” from Xiang et al., were also excluded because of misannotation as previously discussed ^65^. According to published annotation (Epiblast cells from E5-E7, E8-E14, and Carnegie stage 7 were further claimed as “Early Epi”, “Middle Epi” and “Late Epi”, respectively), paired-wise differential expression analysis between lineages were done by using ’roc’ test of FindMarkers function from R package Seurat ^93^. The top 50 up-regulated marker genes with at least average power of more than 0.3 conserved in all comparisons were selected.

## ACKNOWLEDGEMENTS

Images were acquired at the Live Cell imaging Core facility/Nikon Center of Excellence, at the Karolinska Institute, supported by grants from the Swedish Research Council, KI Infrastructure, and Centre for Innovative Medicine. Bioinformatics analyses were performed on resources provided by the Swedish National Infrastructure for Computing (SNIC) at Uppmax server (projects SNIC 2020/15-9, SNIC 2020/6-3).

## SUPPLEMENTARY INFORMATION

**Supplementary Table 1** - Overview of datasets generated for this study and published datasets used

**Supplementary Table 2** - Gene table containing gene names and annotations, promoter H3K4me3, H3K27me3, H2Aub levels, raw RNA-Seq expression values and DESeq2 results generated in this study.

**Extended Data Fig. 1.**
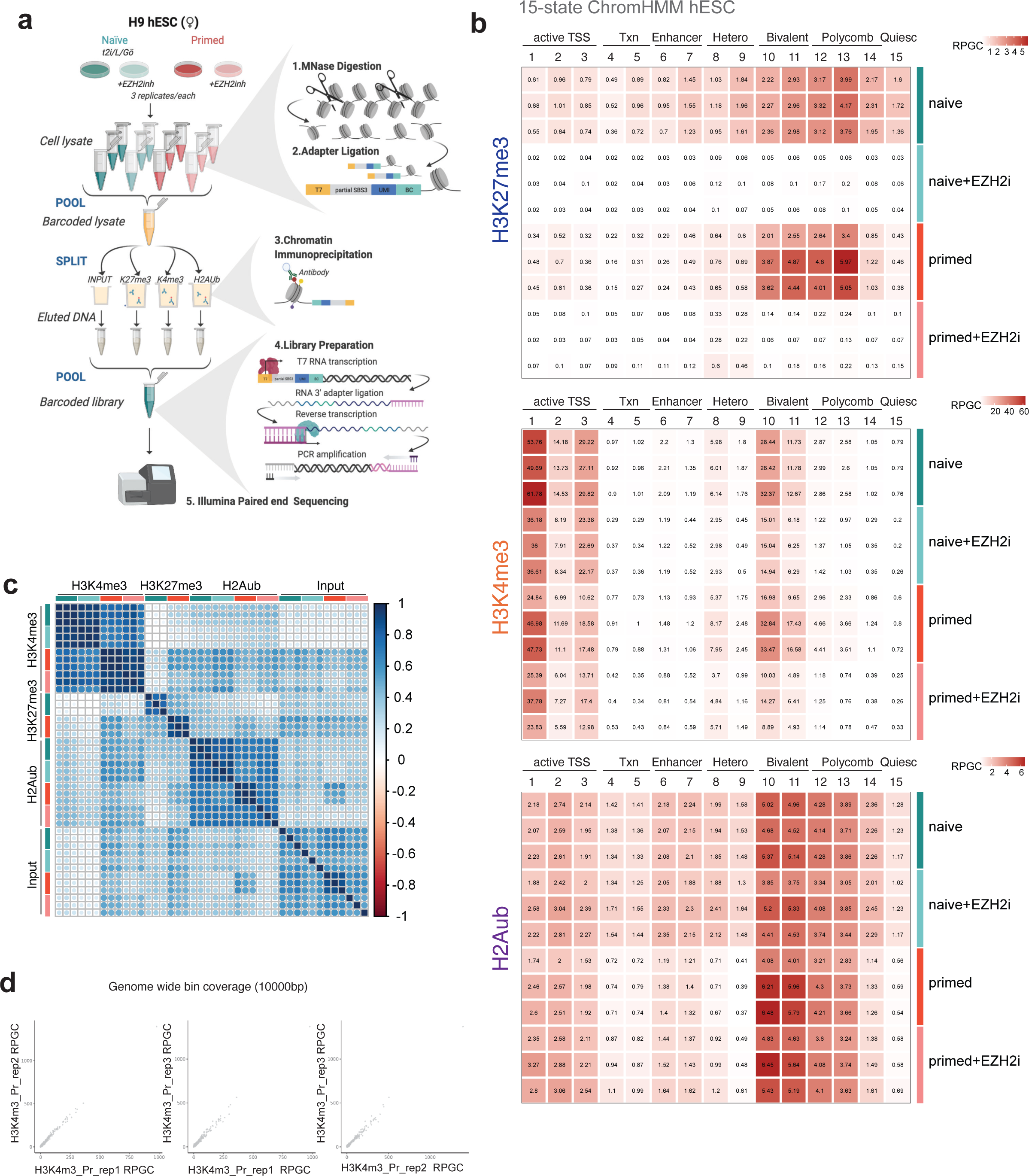
Scheme of MINUTE-ChIP workflow - related to Fig. 1. **a** Scheme of the MINUTE-ChIP workflow. Triplicate cell pellets for each condition were lysed and the chromatin was enzymatically fragmented to mono- and di- nucleosomes and barcoded by ligating on a dsDNA adapter. Samples were then pooled and the barcoded lysate was aliquoted to individual ChIP reactions ( 5% of the ChIP volume was reserved as input material and carried through protocol in a manner similar to the IPs ) with magnetic beads pre-coupled with the respective antibodies. Subsequently beads were washed and IPed DNA was eluted , proteinase K digested and purified. For constructing the final libraries, DNA from each IP was *in vitro* transcribed using the T7 promoter in the adapter that was ligated on in the initial step. The resulting RNA was appended with an RNA 3’ adapter (RA3) allowing for specific paired end sequencing. The RA3 in turn was used to prime the reverse transcription reaction, generating cDNA that was used as a template for the final low-cycle library PCR. At this stage, in addition to the Illumina-compatible sequences, PCR primers also carried a second barcode sequence to serve as an identifier for the IP performed. Finally, libraries are pooled and sequenced on the Illumina platform ^33, 94^. Salient features of this workflow include : The barcode-pool and split strategy which reduces technical variability between samples; the T7 based linear amplification step which requires only one adapter per chromatin fragment to be ligated and the UMI sequence that allows for calling of amplification duplicates. Importantly, the representation of each barcode within the ChIP libraries, compared to the input pool library, yields information for scaling the samples according to their relative global levels for the ChIPed epitope making MINUTE-ChIP a reliable and robust quantitative ChIP-seq method. **b** Histone H3K27me3, H3K4me3 and H2Aub levels by chromatin state. RPGC of individual replicates are shown. **c** Genome-wide correlation (10kb bins, Persson-correlation coefficient) of MINUTE-ChIP replicates, including corresponding inputs. **d** Exemplary genome-wide comparison of 10kb bins as scatter plot

**Extended Data Fig. 2.**
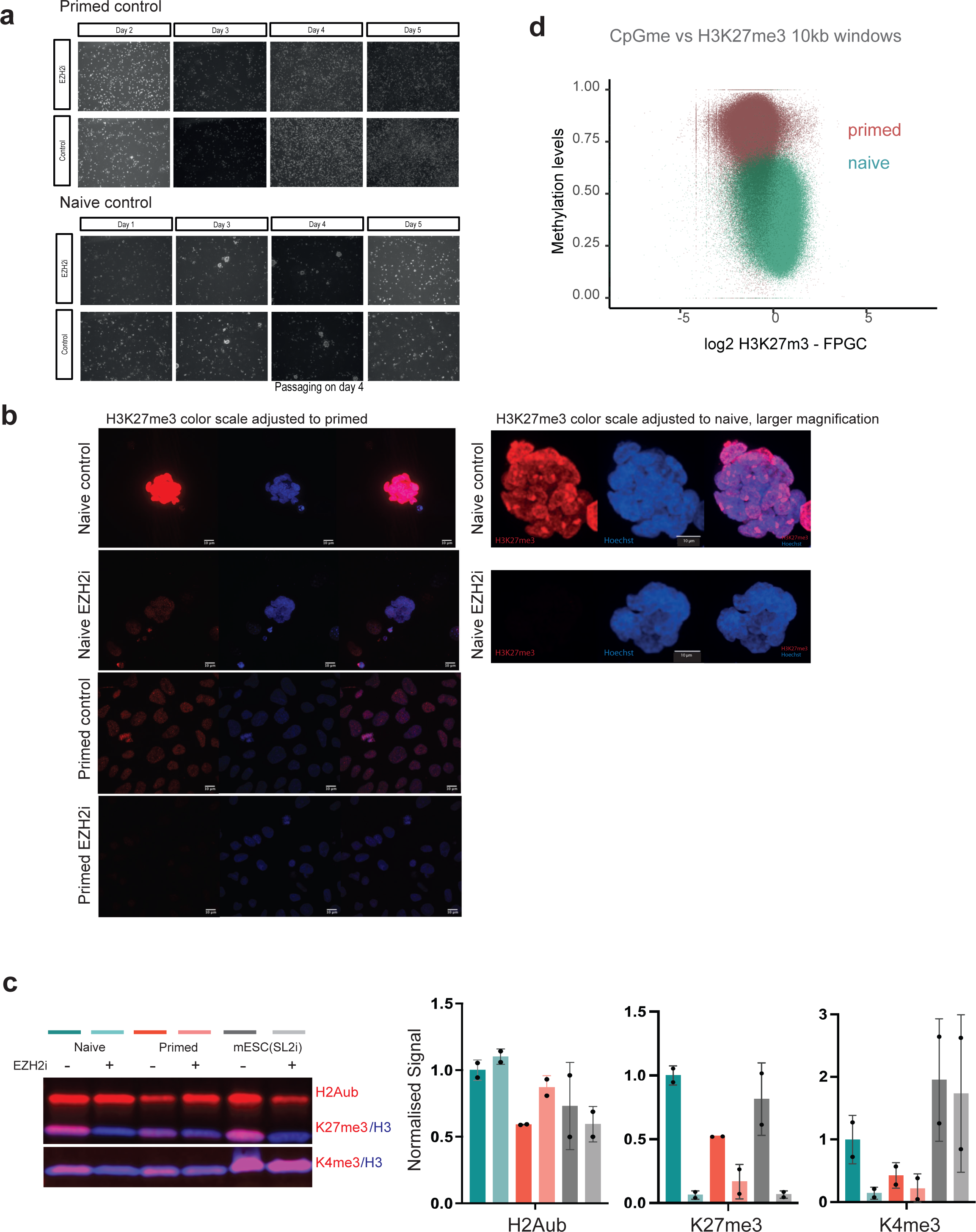
H3K27me3 levels in naïve and primed hESCs - related to Fig. 1. **a** Phase contrast microscopy images of primed and naïve hESC, untreated or treated with EZH2i over 5 days showing similar growth and morphology. **b** Immunofluorescence microscopy showing H3K27me3 staining of naïve and primed hESCs +/- EZH2i treatment, stainings were performed in parallel in the same antibody dilutions and images were acquired with the same gain settings to allow a quantitative comparison. Image analysis yielded a -8fold higher signal in naïve hESC. **c** Protein level H3K4me3, H2Aub and H3K27me3 assayed using two-color IR western blot, in hESCs and WT(J1) mouse ESC (grown in 2i/Serum) with and without EZH2i treatment. Data represented as mean and standard deviation of duplicate samples. Representative image shown. We note that the discrepancy in fold-change comparing naïve and primed hESCs with different quantitative methodologies MINUTE-ChIP, IF, western blots may arise from method-specific threshold sensitivity, dose-response curves and signal saturation levels. MINUTE-ChIP measures H3K27me3-density on the level of nucleosomes, whereas western blot yields a per-histone level quantification. We have previously confirmed that MINUTE-ChIP signal is linear proportional under the dynamic range relevant to the H3K27me3 levels assayed here ^33^. **d** Scatterplot showing weak anticorrelation of H3K27me3 and DNA CpG methylation across 10kb genome wide bins.

**Extended Data Fig. 3.**
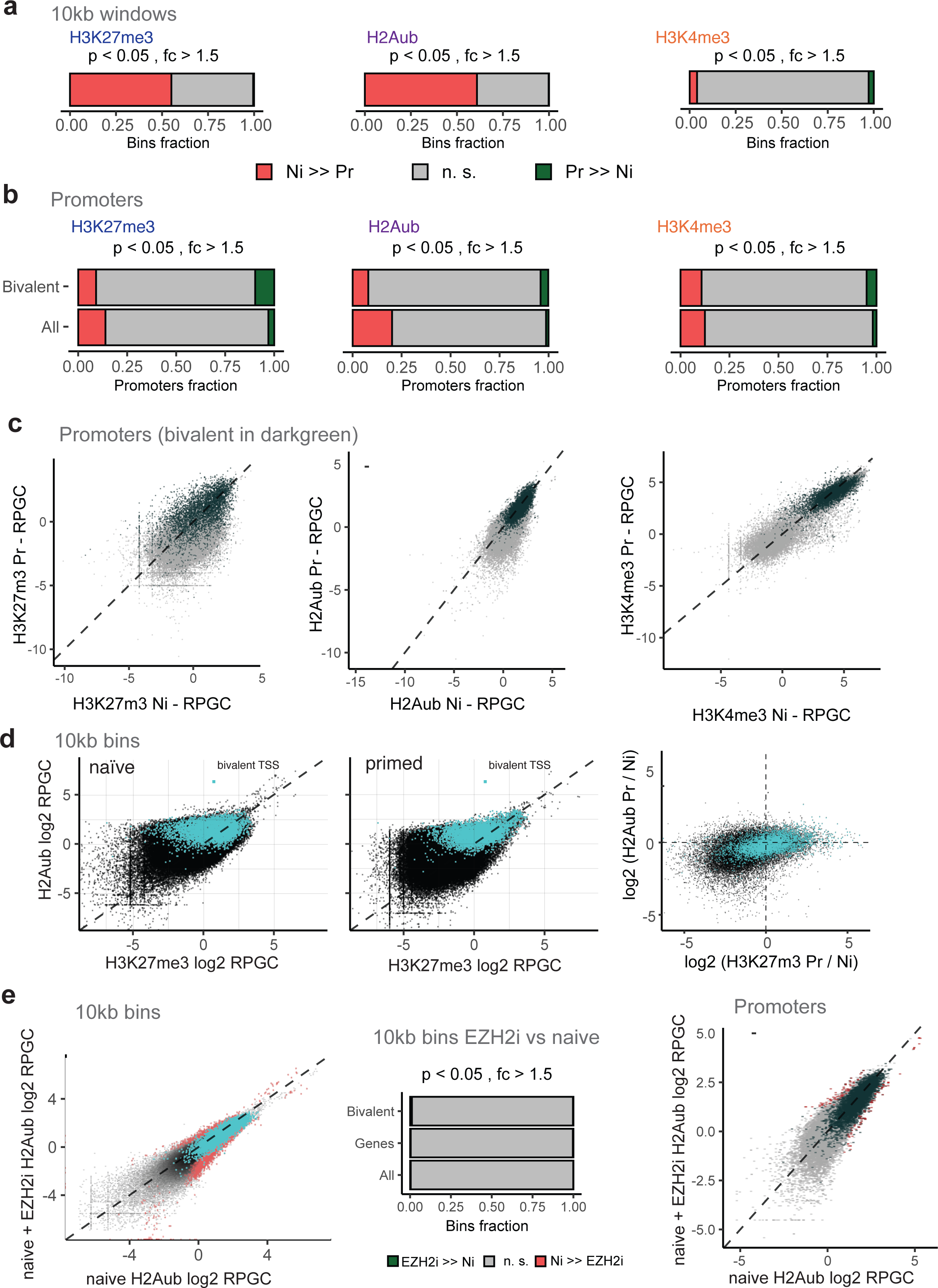
Genome-wide histone modification changes between naïve and primed hESC. related to Fig. 1. **a** Fraction of genome-wide 10kb windows that significantly (DESeq2 p.adj < 0.05 and fold-change > 0.5 from three replicates) gain or lose H3K27me3, H2Aub or H3K4me3 levels between naïve and primed state. **b** Fraction of promoters (all or bivalent ^62^) that significantly (DESeq2 p.adj < 0.05 and fold-change > 0.5 from three replicates) gain or lose H3K27me3, H2Aub or H3K4me3 levels between naïve and primed state. **C** Scatter plots showing H3K27me3, H2Aub or H3K4me3 levels (log2-transformed RPGC) at promoters in naïve verus primed hESC. Bivalent genes are highlighted in black. **d** Scatter plots showing the relation of H3K27me3 and H2Aub across genome-wide 10kb bins in naïve and primed hESC, as well as the correlation of changes. 10kb bins overlapping with bivalent TSS are highlighted in turquoise **e** Scatter plot showing the relation of H2Aub across genome-wide 10kb bins in naïve vs. EZH2i-treated naïve hESC. Significant bins are highlighted in red. 10kb bins overlapping with bivalent promoters are highlighted in turquoise. Fraction of 10kb bins significantly changing (DESeq2 p.adj < 0.05 and fold-change > 0.5 from three replicates) are given for all 10kb bins, 10kb bins overlapping with promoters, and 10kb bins overlapping with bivalent promoters. Scatter plot showing the relation of H2Aub across promoters in naïve vs. EZH2i-treated naïve hESC. Significant bins are highlighted in red. Bivalent promoters are highlighted in black.

**Extended Data Fig. 4.**
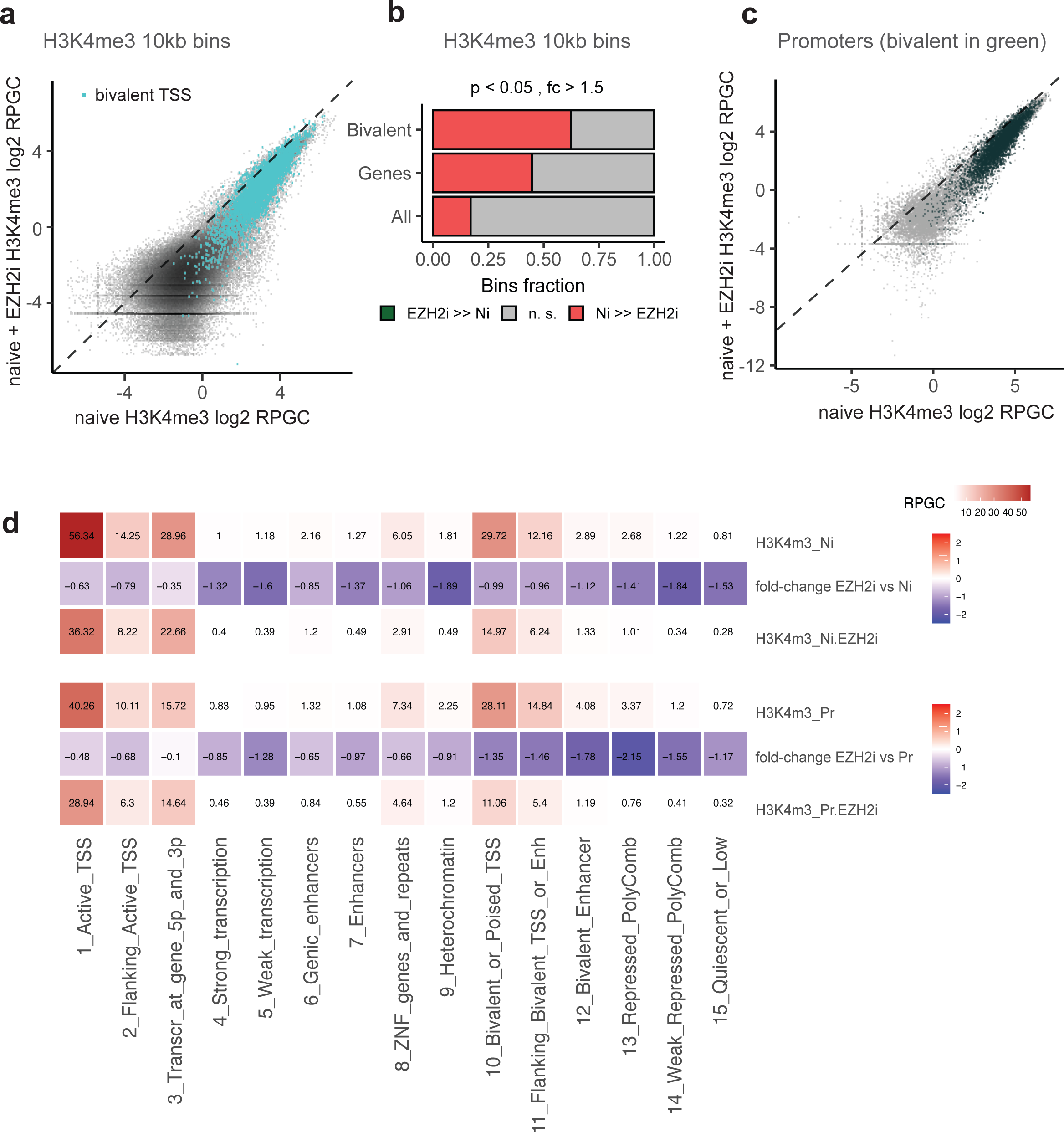
Genome-wide changes in H3K4me3 in response to EZH2i treatment. related to Fig. 1. **a** Scatter plot showing the levels of H3K4me3 across genome-wide promoters in naïve vs. EZH2i-treated naïve hESC. Significant bins are highlighted in red. Bivalent promoters are highlighted in black. H3K4me3 levels are reduced relatively evenly by -2-fold after EZH2i treatment. **b** Fraction of 10kb bins significantly changing (DESeq2 p.adj < 0.05 and fold-change > 0.5 from three replicates) are given for all 10kb bins, 10kb bins overlapping with promoters, and 10kb bins overlapping with bivalent promoters. **c** Scatter plot showing the levels of H3K4me3 across promoters in naïve vs. EZH2i-treated naïve hESC. Significant bins are highlighted in red. Bivalent promoters are highlighted in black. **d** Histone H3K4me3 levels by chromatin state, naïve and primed hESCs +/- EZH2i treatment. RPGC of combined replicates are shown. Two additional rows show log2-fold gains or losses comparing EZH2i-treated and untreated cells. While H3K4me3 reduced by -35% at active TSS, the average reduction at bivalent promoters was 50%, and background levels were reduced by 65-75%. The EZH2 inhibitor used here, EPZ-6438, does not affect H3K4me3 levels in other cell types ^34^ and mESC (Extended Data Fig. 2C), thus the reduction in H3K4me3 levels appears to be a specific response to H3K27me3 depletion in naïve and primed hESC.

**Extended Data Fig. 5.**
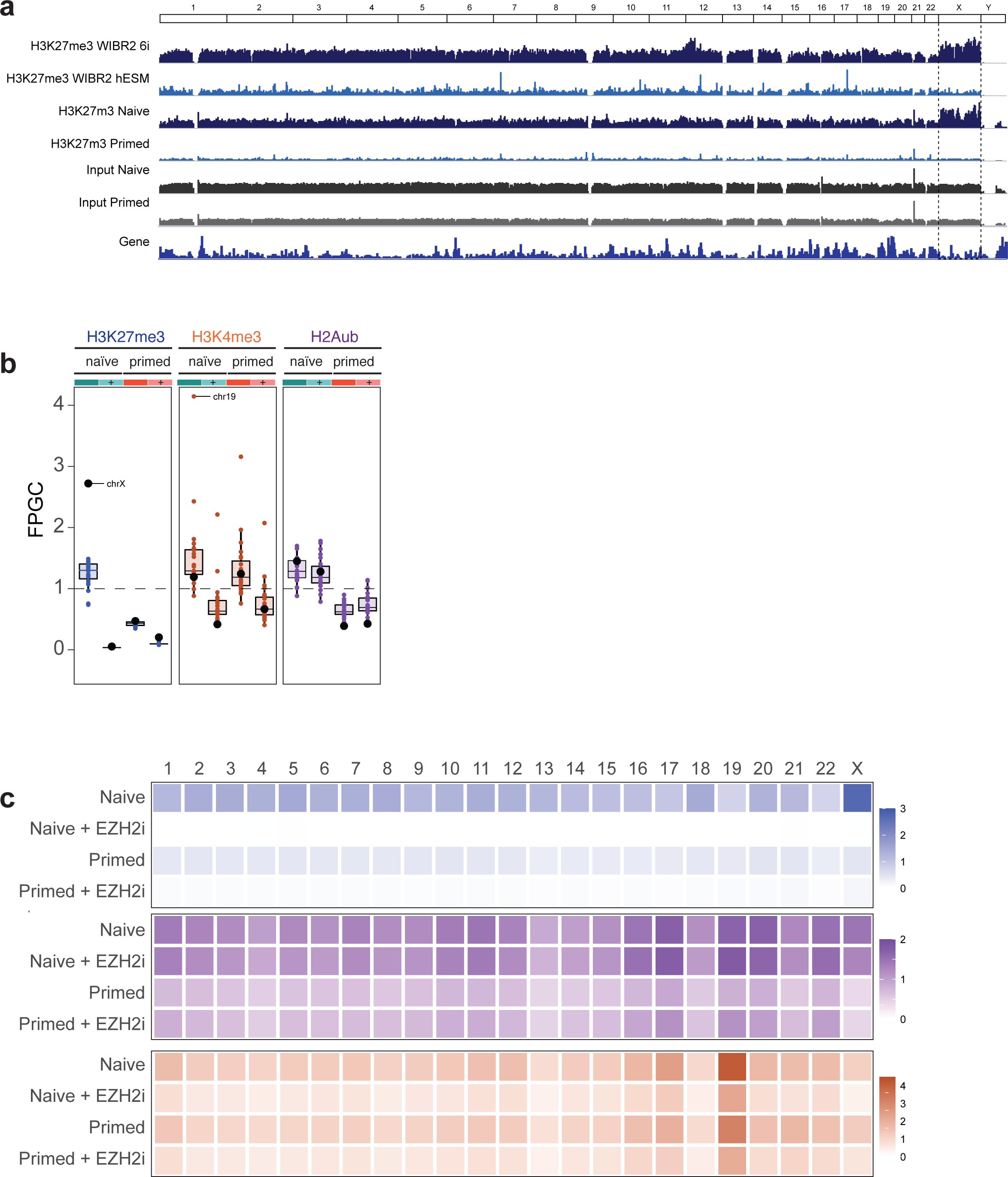
High H3K27me3 density on naïve X chromosomes - related to Fig. 2. **a** Representative Genome browser overview showing H3K27me3 density across all chromosomes. Published non-quantitative H3K27me3 ChIP (top, ^17^) and MINUTE-ChIP H3K27me3 tracks (combined replicates), as well as MINUTE-ChIP input tracks are shown. **b** Chromosome average enrichment of H3K4me3, H3K27me3 and H2Aub in naïve and primed hESC, cultured with or with EZH2i for 7 days. RPGC of combined replicates is shown. **c** Heatmap showing per-chromosome average H3K27me3, H2Aub and H3K4me3 signal in naïve and primed hESC, cultured with or with EZH2i for 7 days.

**Extended Data Fig. 6.**
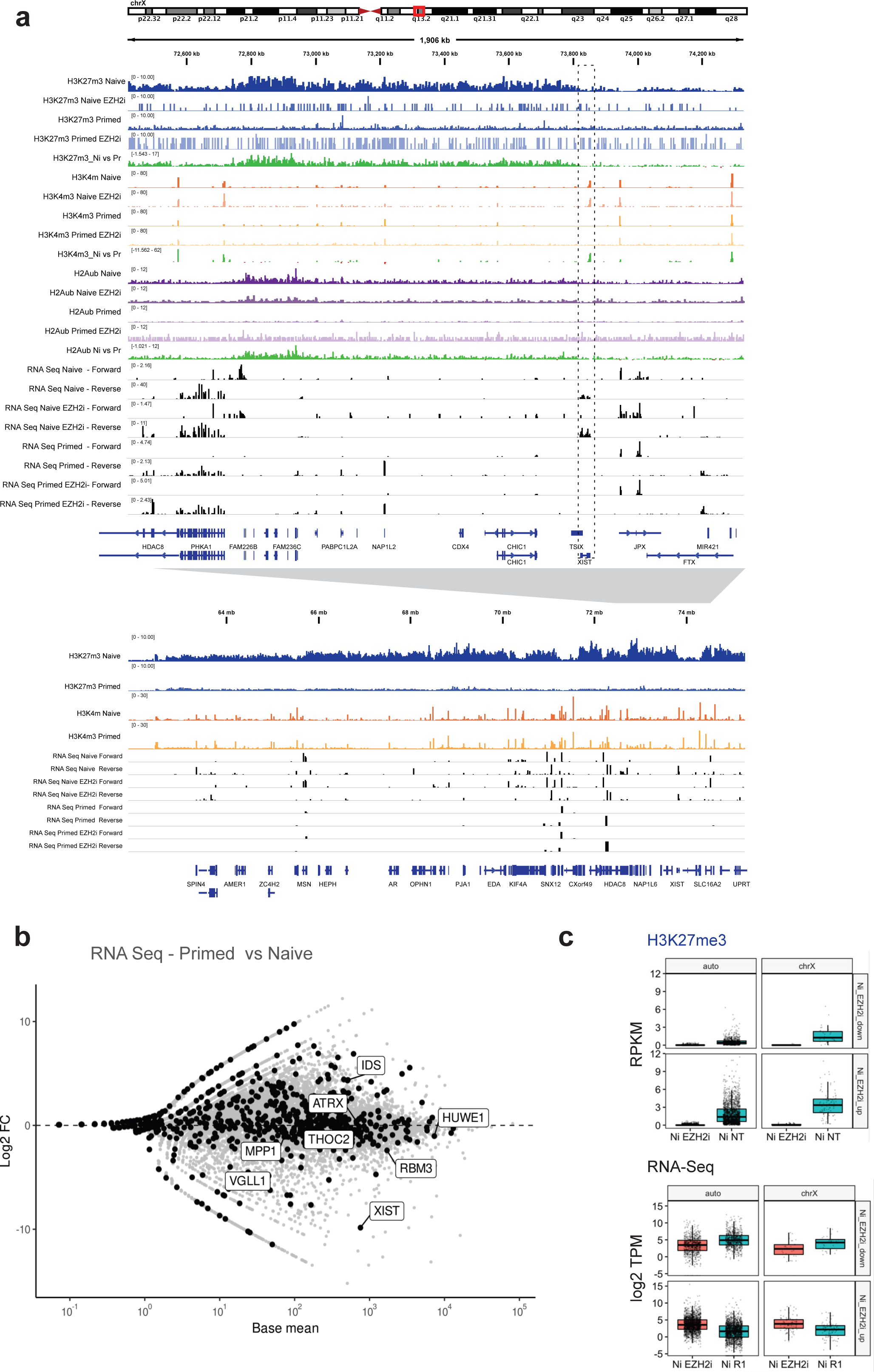
Comparison of epigenetic and transcriptional status of the X chromosome in naïve and primed hESCs - related to Fig. 2. **a** Genome browser view of XIST neighbourhood on X chromosome at megabase resolution. Shown is the combined signal of three replicates for H3K27me3, H3K4me3, H2Aub as well as gain/loss tracks comparing naïve and primed signal, and stranded RNA-Seq signal. Tracks from the same histone modification are shown on the same RPGC scale. RNA-Seq expression is shown on the same TPM scale. Strong H3K27me3 accumulation in large blocks on naïve X chromosome is evident, with the exception of some islands that typically harbor genes active in naïve state, such as the XIST locus, which do not accumulate H3K27me3. **b** MA-plot of RNA-Seq data (Base mean and log2 fold-change as calculated with DESeq2 from triplicates) comparing naïve and primed hESC. Genes on the X chromosome are highlighted in black with selected annotations. **c** Boxplots showing H3K27me3 levels at promoters on the X chromosome or autosomes in naïve hESCs +/- EZH2i treatment (top). Two groups of promoters are plotted: those of genes upregulated with EZH2i treatment in naïve hESCs and those downregulated. Boxplots showing expression levels (log2-transformed TPM) at genes on the X chromosome or autosomes in naïve and primed hESCs +/- EZH2i treatment (bottom).

**Extended Data Fig. 7.**
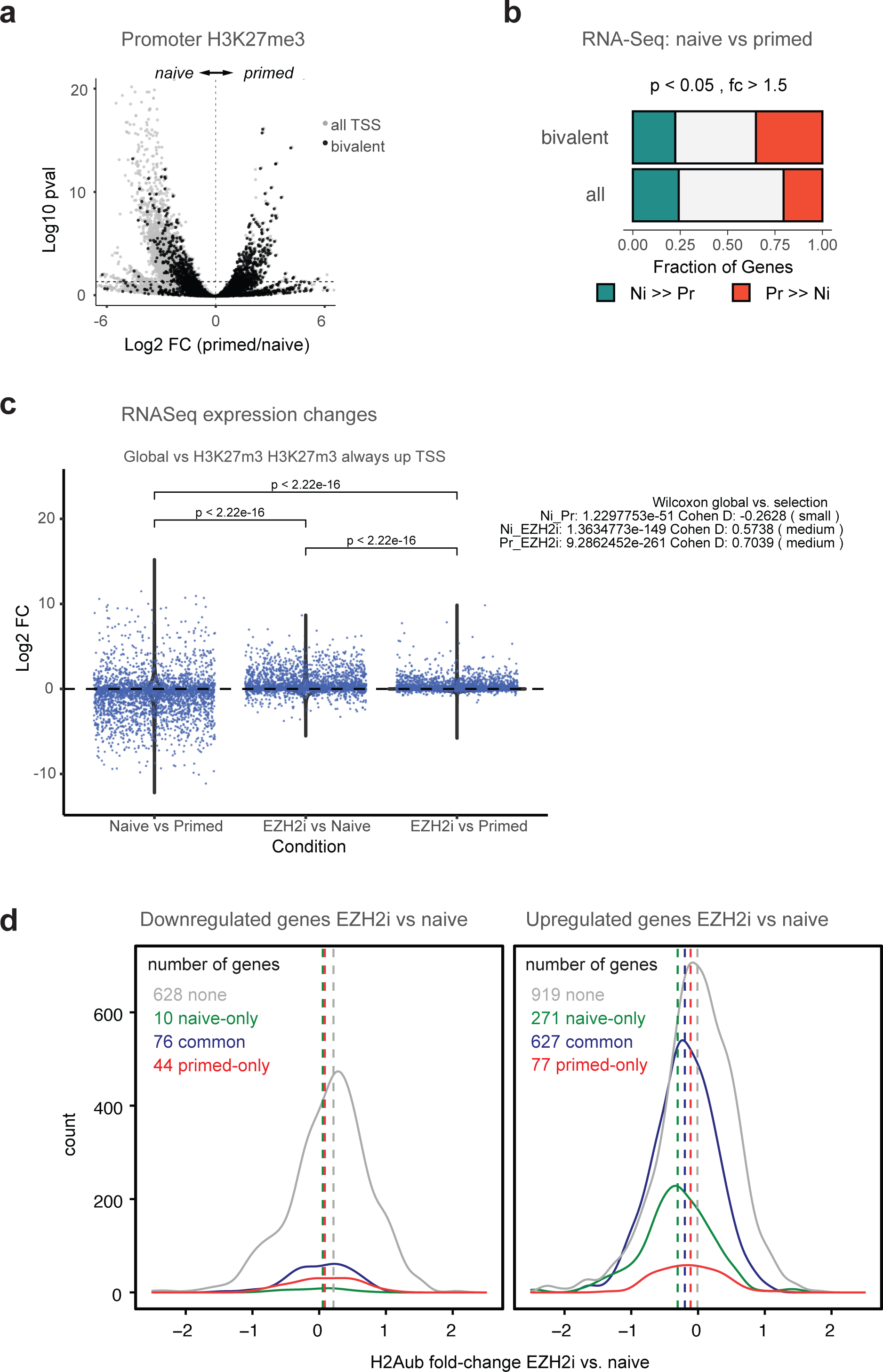
Context specific changes in bivalency between pluripotent states and response to H3K27me3 depletion - related to Fig. 3. **a** Volcano plot (DESeq2 based on three replicates) comparing promoter H3K27me3 levels between naïve and primed hESC. Bivalent promoters ^62^ are highlighted in black. **b** Fraction of significantly up- or downregulated genes (DESeq2 p.adj < 0.05, fold-change > 1.5 based on three replicates) comparing naïve and primed hESCs amongst all genes, or annotated bivalent genes from primed hESCs ^62^. Overall, a similar fraction of genes was up- and downregulated significantly comparing naïve vs primed hESCs (-25% each), but amongst bivalent genes, 24% were downregulated and 35% upregulated in primed hESCs **c** State-specific transcriptional response to global H3K27me3 depletion in shared bivalent genes. RNA-Seq changes (log2 fold-change from DESeq2 analysis from three replicates each) are plotted, comparing naïve and primed condition as well as EZH2i treatment and the respective control condition. The distribution of fold-changes of all genes is shown as violin plots, whereas the class-specific genes are shown as jitter points. Significance of pairwise Student’s t test is given. The mean fold-changes of the class-specific group was compared to the mean fold-change of all genes using Students t-test and Cohen’s d. **d** Density plot of fold-changes of H2Aub levels following H3K27me3 depletion in hESC. Only genes that were derepressed upon EZH2i-treatment (DESeq2 p.adj < 0.05, fold-change > 1.5 based on three replicates) were included in the analysis. The different classes of bivalent promoters as defined in Fig. 3B are compared to H3K27me3-devoid promoters.

**Extended Data Fig. 8.**
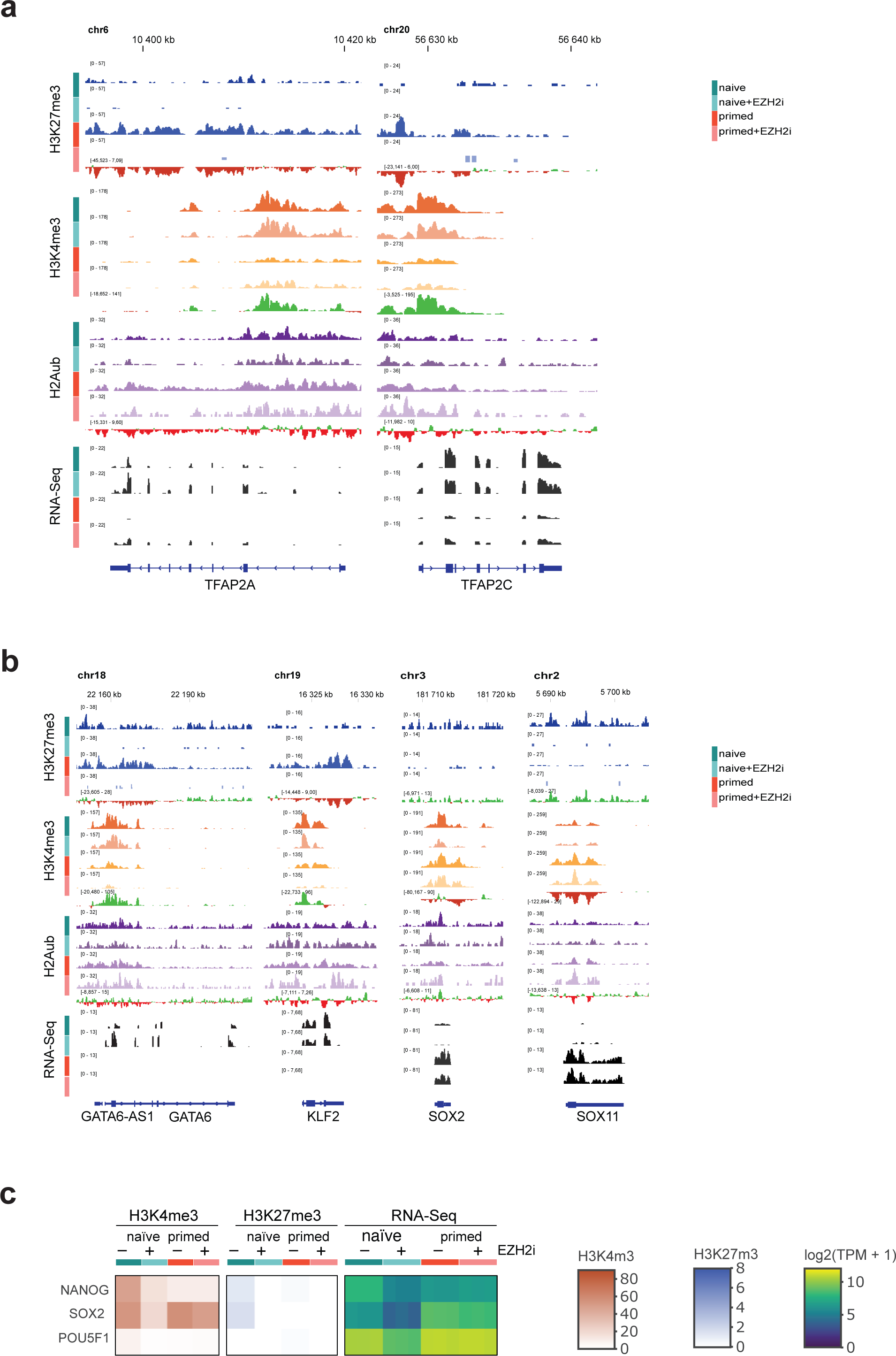
Genome browser examples of naïve and primed-specific pluripotency markers - related to Fig. 4. **a** Genome browser view of TFAP2A and TFAP2C transcription factors in naïve and primed hESCs +/- 7d EZH2i treatment. Shown is the combined signal of three replicates for H3K27me3, H3K4me3, H2Aub as well as gain/loss tracks comparing naïve and primed signal, and stranded RNA-Seq signal. Tracks from the same histone modification are shown on the same RPGC scale. RNA-Seq expression is shown on the same TPM scale. **b** Genome browser view of naïve or primes-specific pluripotency marker genes, in naïve and primed hESCs +/- 7d EZH2i treatment. Shown is the combined signal of three replicates for H3K27me3, H3K4me3, H2Aub as well as gain/loss tracks comparing naïve and primed signal, and stranded RNA-Seq signal. Tracks from the same histone modification are shown on the same RPGC scale. RNA-Seq expression is shown on the same TPM scale. **c** Heatmap showing RNA-Seq expression levels (log2-transformed TPM) of core pluripotency markers in naïve and primed hESCs +/- 7d EZH2i treatment, as well as the H3K4me3 and H3K27me3 levels (RPGC) at their respective promoter. RPGC from combined replicates are used for H3K4me3 and H3K27me3, whereas the three individual replicate TPM values are plotted for RNA-Seq data.

**Extended Data Fig. 9.**
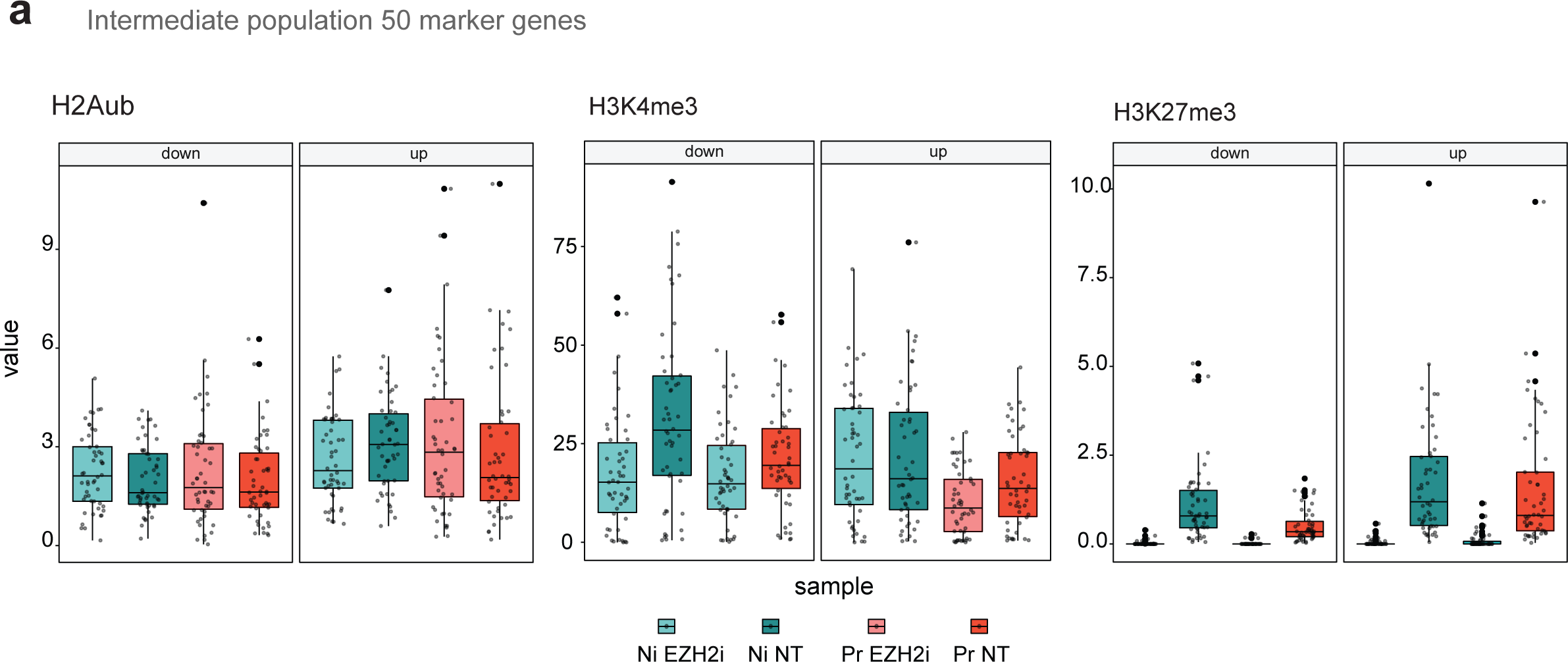
Chromatin features of genes uniquely up- or downregulated in the ’intermediate’ naïve population - related to Fig. 4. Boxplots showing H3K27me3, H3K4me3 and H2Aub levels (RPGC) of naïve and primed cells +/- 7d EZH2i treatment, for marker genes of an ’intermediate’ population present at -2% abundance in naïve cultures as defined in Messmer et. al. 2019 ^64^. The 50 top up- and down-regulated genes reported for the ’intermediate’ population were selected.

**Extended Data Fig. 10.**
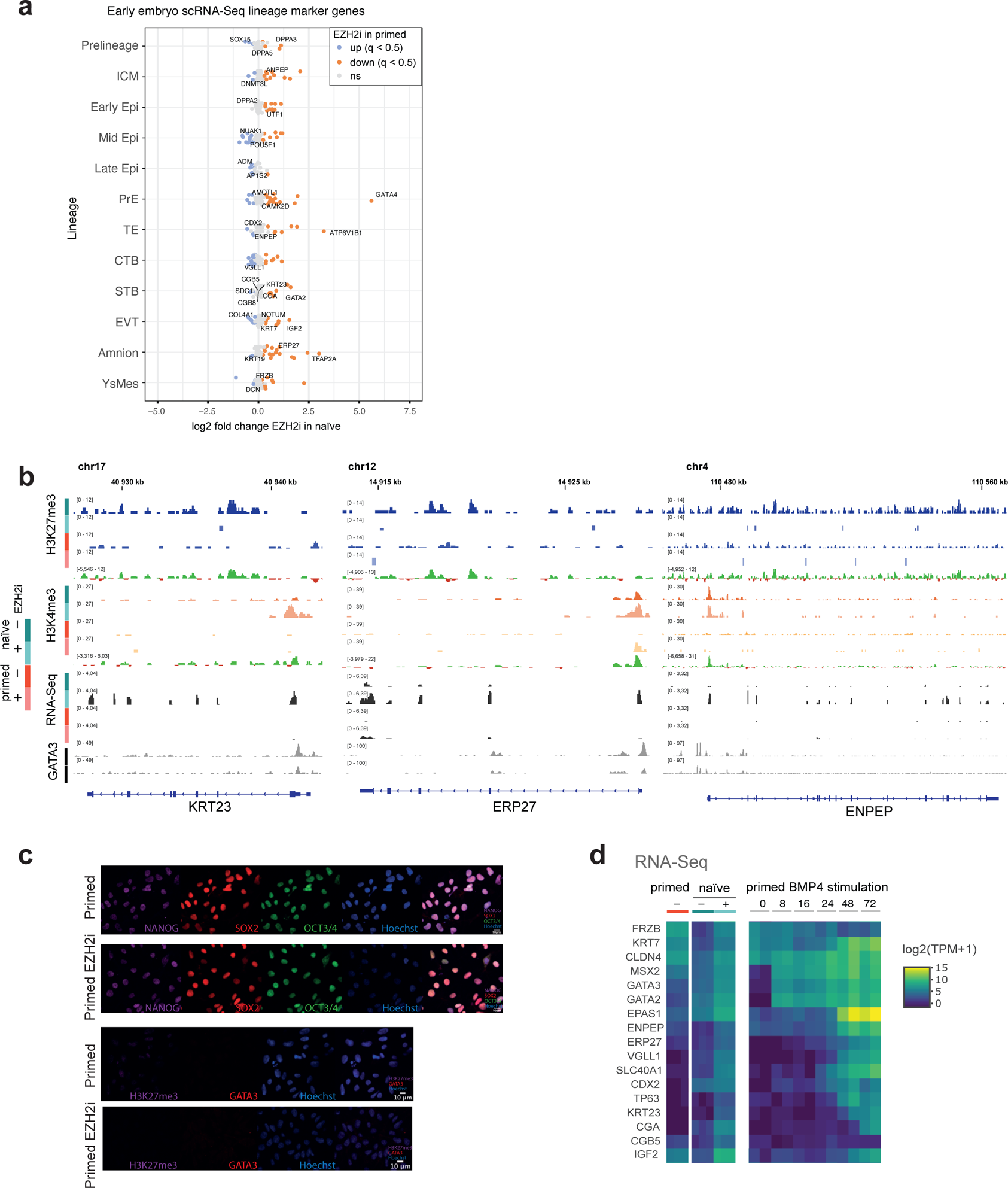
Loss of H3K27me3 in naïve activates trophectoderm gene expression programs - related to Fig. 5. **a** Barchart showing sets of marker genes for different cell fates as determined by single-cell transcriptomics. Markers are grouped into pre-lineage, inner cell mass (ICM), epiblast, primitive endooderm, (PrimEndo), trophectoderm (TE) cytotrophoblast (CTB), syncytiotrophoblast (STB), extravillous trophoblast (EVT), amnion and yolk sac mesoderm (YsMes) and primitive streak (PriS). For each set of marker genes, the number of genes that significantly increase or decrease their expression upon EZH2i treatment in naïve, primed or both states are indicated. **b** Genome browser examples of selected trophectoderm lineage markers with H3K27me3-less promoters. Shown is the combined signal of three replicates for H3K27me3, H3K4me3, H2Aub, RNA-Seq, as well as gain/loss tracks comparing naïve and primed signal. Tracks from the same histone modification are shown on the same RPGC scale. RNA-Seq expression is shown on the same TPM scale. **c** Immunofluorescence microscopy showing H3K27me3, OXT3/4, SOX2, NANOG, GATA3 staining of naïve and primed hESCs +/- EZH2i treatment. **d** Heatmap comparing trophectoderm marker gene expression in our primed, naïve, naïve+EZH2i datasets with a time course experiment reprogramming primed hESCs with BMP4 ^9^. RNA-Seq expression is shown on the same TPM scale, triplicates are given for our dataset and duplicates for the published datasets.

**Extended Data Fig. 11.**
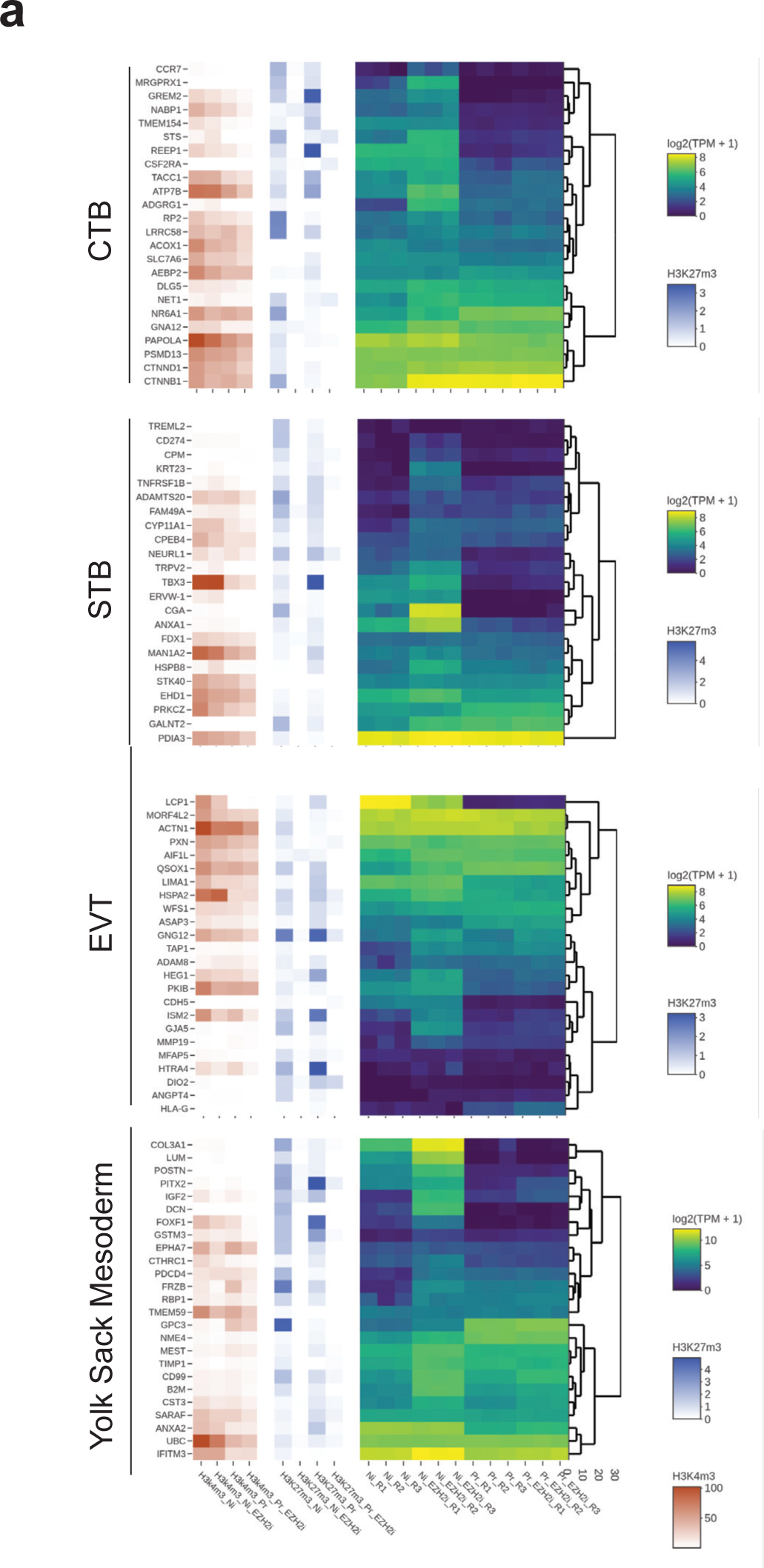
Loss of H3K27me3 in naïve activates placental genes - related to Fig. 5. Heatmaps showing RNA-Seq expression levels (log2-transformed TPM) of core pluripotency markers in naïve and primed hESCs +/- 7d EZH2i treatment, as well as the H3K4me3 and H3K27me3 levels (RPGC) at their respective promoter. RPGC from combined replicates are used for H3K4me3 and H3K27me3, whereas the three individual replicate TPM values are plotted for RNA-Seq data.

